# Co-evolving wing spots and mating displays are genetically separable traits in *Drosophila*

**DOI:** 10.1101/869016

**Authors:** Jonathan H. Massey, Gavin R. Rice, Anggun Firdaus, Chi-Yang Chen, Shu-Dan Yeh, David L. Stern, Patricia J. Wittkopp

## Abstract

The evolution of sexual traits often involves correlated changes in morphology and behavior. For example, in Drosophila, divergent mating displays are often accompanied by divergent pigment patterns. To better understand how such traits co-evolve, we investigated the genetic basis of correlated divergence in wing pigmentation and mating display between the sibling species *Drosophila elegans* and *D. gunungcola*. *Drosophila elegans* males have an area of black pigment on their wings known as a wing spot and appear to display this spot to females by extending their wings laterally during courtship. By contrast, *D. gunungcola* lacks both of these traits. Using Multiplexed Shotgun Genotyping (MSG), we identified a ∼440 kb region on the X chromosome that behaves like a genetic switch controlling the presence or absence of male-specific wing spots. This region includes the candidate gene *optomotor-blind* (*omb*), which plays a critical role in patterning the *Drosophila* wing. The genetic basis of divergent wing display is more complex, with at least two loci on the X chromosome and two loci on autosomes contributing to its evolution. Introgressing the X-linked region affecting wing spot development from *D. gunungcola* into *D. elegans* reduced pigmentation in the wing spots but did not affect the wing display, indicating that these are genetically separable traits. Consistent with this observation, broader sampling of wild *D. gunungcola* populations confirmed the wing spot and wing display are evolving independently: some *D. gunungcola* males preformed wing displays similar to *D. elegans* despite lacking wing spots. These data suggest that correlated selection pressures rather than physical linkage or pleiotropy are responsible for the coevolution of these morphological and behavioral traits. They also suggest that the change in morphology evolved prior to the change in behavior.

## Introduction

Animals often use colorful morphological structures to communicate with prospective mates during courtship (McKinnon and Pierotti, 2010). In vertebrates and invertebrates, pigmented bodies or wings often evolve together with specific components of courtship behavior that animals use to display their colorful anatomy (Loxton, 1979; Endler, 1991; Sinervo et al., 2000; White et al., 2015). These correlated differences evolve both within and between populations, frequently distinguishing males from females or closely related species (Gray and McKinnon, 2007; McKinnon and Pierotti, 2010). In the handful of case studies examining the genetic basis of such co-evolving traits, linkage mapping and genome-wide association studies (GWAS) have shown that loci affecting pigmentation patterning tend to co-localize with loci affecting variation in mating behaviors (Lindholm and Breden, 2002; Kronforst *et al*., 2006; Yeh *et al.*, 2006; Thomas *et al*., 2008; Kupper *et al*., 2016; Lamichhaney *et al*., 2016; Merrill *et al*., 2019; reviewed in McKinnon and Pierotti, 2010). That is, physical linkage of genetic variants underlies phenotypic correlations between mating behavior and pigmentation. Interestingly, these loci also tend to explain much of the variation observed for both traits (e.g., Kronforst *et al*., 2006; Kupper *et al*., 2016; Lamichhaney *et al*., 2016). A key challenge is determining how frequently these patterns of genomic architecture underlie correlated evolution and whether a single pleiotropic locus or separate linked loci are involved.

Disentangling whether pleiotropic or physically linked loci underlie patterns of correlated evolution between pigmentation and mating behavior is important for understanding how natural selection generates differences between sexes and species. If two beneficial traits are genetically correlated due to separate, physically linked loci, theory predicts that natural or sexual selection (e.g., through predation or female choice) will act to minimize recombination between the causal loci (Charlesworth and Charlesworth, 1976). It has been hypothesized that one solution to this problem might involve the evolution of chromosomal inversions that suppress recombination between two or more linked loci (Kirkpatrick and Barton, 2006). Alternatively, mutations at a single pleiotropic gene could cause correlated components of pigmentation and mating behavior to evolve simultaneously, although it is not likely, mechanistically, that a single mutation with generate adaptive changes in both pigmentation and behavior. Distinguishing between these genetic modes of phenotypic evolution requires, in part, high-resolution mapping of correlated traits.

In the Oriental *Drosophila melanogaster* species group, male-specific wing spots are phylogenetically correlated with mating displays (Kopp and True, 2002; Figure 1A). Species with wing spots perform elaborate wing display dances during courtship, extending their wings laterally, turning their dorsal wing surfaces toward the female, and waving them up and down; species without wing spots lack display behavior (Kopp and True, 2002, Figure 1A,B). Correlated gains and losses of both traits have evolved repeatedly (Kopp and True, 2002, Figure 1A). For example, in *D. elegans* and *D. gunungcola*, sibling species from this group that are estimated to have diverged 2-2.8 million years ago (Prud’homme *et al*., 2006), *D. elegans* (Bock and Wheeler, 1972) males possess wing spots and perform wing displays, whereas *D. gunungcola* (Sultana *et al*., 1999) males lack both traits (Kopp and True, 2002; Prud’homme *et al*., 2006; Yeh *et al*., 2006; Figure 1B; Video 1; Video 2). Previously, Yeh *et al*., (2006) and Yeh and True (2014) discovered that *D. elegans* and *D. gunungcola* can generate fertile F_1_ hybrid female offspring in the lab and they performed interspecific crosses to study the genetic basis of wing spot and wing display divergence. Through quantitative trait locus (QTL) mapping, they showed that evolution of linked loci on the X chromosome contributed to divergence in both traits (Yeh *et al.*, 2006; Yeh and True, 2014). One QTL explaining wing spot size variation was linked to the pigmentation gene *yellow*, supporting the hypothesis that *yellow cis*-regulatory divergence contributes to wing pigmentation evolution (Wittkopp *et al*., 2002a; Gompel *el al*., 2005; Prud’homme *et al*., 2006). It remained unclear, however, whether the same or different loci on the X chromosome underlie correlated differences in wing spot and wing display between these species.

**Figure 1.**
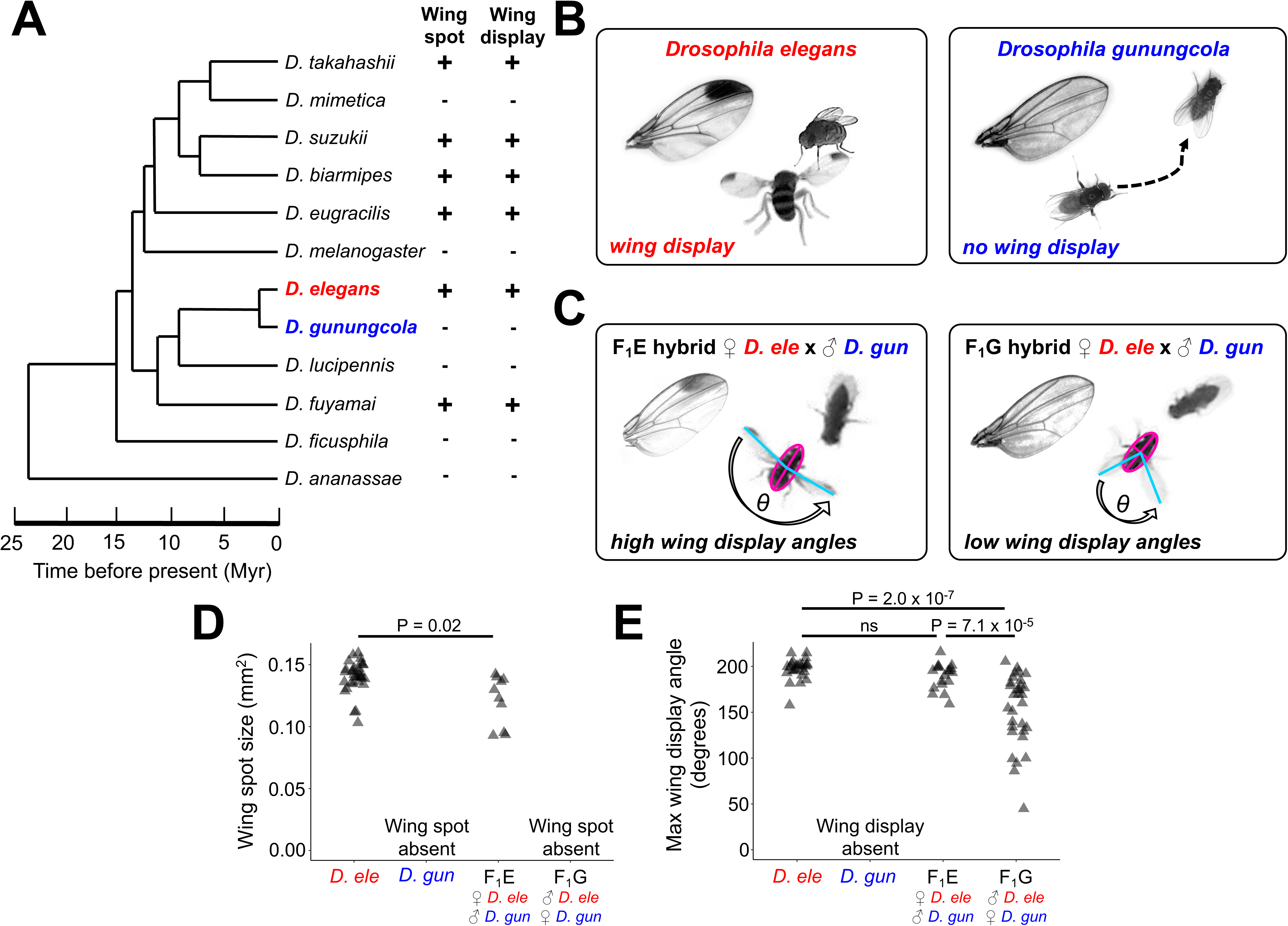
Wing pigmentation and wing display behavior in *D. elegans*, *D. gunguncola*, and F_1_ hybrids. (A) Phylogeny of the “Oriental” *Drosophila melanogaster* species group adapted from Kopp and True (2002) and Prud’homme *et al*. (2006). Plus (+) signs indicate species possess wing spots and/or wing displays, and minus (-) signs indicates wing spots and/or wing displays are absent. (B) Males in *D. elegans* (left) possess wing spots and perform bilateral wing display behaviors in front of females during courtship (Video 1). Wing spots and wing displays are absent in *D. gunungcola* males (right) (Video 2). (C) F_1_ hybrid males inheriting their X chromosome from *D. elegans* mothers (F_1_E, left) possess wing spots and perform wing display behavior like *D. elegans* (Video 3). F_1_ hybrid males inheriting their X chromosome from *D. gunungcola* mothers (F_1_G, right) are spotless and perform wing displays with low bilateral wing angles (Video 4). (D) Quantification of wing spot size (see Methods) in male *D. elegans* and F_1_E. Wing spots are larger in *D. elegans* than F_1_E (Student’s t-test; t = −2.8057; df = 11.43; P = 0.017; two-tailed). (E) Quantification of maximum bilateral wing display angles during courtship (see Methods) in male *D. elegans* and F_1_ hybrids. F_1_G hybrids showed lower maximum wing display angles than *D. elegans* and F_1_E hybrids (One-way ANOVA: F_2,71_ = 20.92; P < 7.18 x 10^-8^; post-hoc Tukey HSD was significant between *D. elegans* and F_1_G: P < 2.0 x 10^-7^ and between F_1_E and F_1_G: P < 7.1 x 10^-5^). Gray triangles represent individual replicates.

To distinguish between these possibilities, we re-examined the genetic basis of wing spots and wing display divergence between *D. elegans* and *D. gunungcola.* Specifically, we (1) generated recombinant backcross progeny segregating for both traits, (2) assembled chromosome-length scaffolds of *D. elegans,* (3) used Multiplexed Shotgun Genotyping (MSG) (Andolfatto *et al*., 2011) to estimate recombination crossover positions across the genome, (4) generated quantitative measures of both wing spots and wing display behavior to estimate the effect size of loci contributing to divergence, and (5) generated advanced, recombinant introgressions on the X chromosome in an attempt to separate quantitative trait loci (QTL) underlying wing spots and wing display behavior. These experiments showed that a single locus on the X chromosome behaves like a genetic switch for wing spot divergence; however spotless males inheriting introgressions of this region from *D. gunungcola* in a *D. elegans* genetic background performed wing displays like *D. elegans* males, indicating that the two traits are genetically separable. These findings suggest that wing spot and wing display behavior might have originally diverged independently. Consistent with this hypothesis, newly collected *D. gunungcola* strains from Indonesia appear to completely lack wing spots but retain the ability to perform wing displays. This observation suggests that the loss of wing spots occurred prior to the loss of wing display in the reference strain of *D. gunungcola* used in this study and in prior work.

## Materials and Methods

### Fly stocks

The *D. elegans HK* (Hong Kong) and *D. gunungcola SK* (Sukarami) lines used in this study were a gift from John True (Stony Brook University). Species stocks were kept on a 12 h light-dark cycle at 23°C on a University of Michigan “R food” diet containing molasses (http://lab-express.com/flyfoodsupplies.htm#rfood) (Wirtz and Semey, 1982). Maintaining these species on R food at high densities (50-100 flies per vial) allowed for the parental population to build up to thousands of flies to collect hundreds of virgins for interspecific crosses (see below). Neither *D. elegans* nor *D. gunungcola* pupate on the sides of the vial, so adults were flipped out when 3^rd^ instar L3 larvae developed and Fisherbrand filter paper (cat# 09-790-2A) was added to the food to create pupation space.

### Generating hybrid progeny

Virgin males and females of *D. elegans* and *D. gunungcola* were isolated upon eclosion and stored in groups of ten for one week on University of Michigan “M food”, which is the standard cornmeal diet from the Bloomington Drosophila Stock Center (https://bdsc.indiana.edu/information/recipes/bloomfood.html) with 20% higher agar content. Virgin males from *D. elegans* were crossed to virgin females from *D. gunungcola*, and virgin males from *D. gunungcola* were crossed to virgin females from *D. elegans* in groups of ten males and ten females to generate fertile F_1_ female and sterile F_1_ male hybrids. These crosses took ∼3-4 weeks to produce hybrid progeny. The switch from R food to M food for interspecific crosses was necessary, because R food tended to accumulate condensation and bacterial growth much faster than M food when few flies occupied a vial. Since crossing *D. elegans* and *D. gunungcola* to generate F_1_ hybrids tends to take several more weeks than within species crosses, the switch to M food diet allowed for maximum breeding time and the development of dozens of hybrid progeny. Once hybrid females eclosed from both interspecific cross directions, they were pooled into the same vial and aged for ten days. We did not keep track of F_1_ hybrid female maternity, because previous work (Yeh and True, 2014) found no effect of F_1_ hybrid maternity on trait means for wing spots and wing display in backcross populations. Multiple high-density groups of ∼60 F_1_ hybrid females were then backcrossed to ∼60 virgin male *D. elegans* flies in individual vials on M food diet to create the *D. elegans* backcross recombinant population (724 individuals). To create the *D. gunungcola* backcross recombinant population (241 individuals), groups of ∼60 F_1_ hybrid females were backcrossed to ∼60 virgin male *D. gunungcola* flies in individual vials on M food diet; this backcross was less successful at producing recombinant progeny than the *D. elegans* backcross direction.

### Behavioral assays

Virgin *D. elegans* females were isolated upon eclosion, aged 10-20 days, and stored in groups of 30-40 for courtship assays. F_1_ hybrid and recombinant backcross males were isolated individually in M food vials using CO_2_ upon eclosion for at least 5 days before each courtship assay. For each assay, a single individual male was gently aspirated into a custom built 70 mm diameter bowl arena that matches the specifications in Simon and Dickinson (2010). Next, a single virgin *D. elegans* female was aspirated into the chamber and videotaped for the next 20 min, using a Canon VIXIA HF R500 camcorder mounted to Manfrotto (MKCOMPACTACN-BK) aluminum tripods. Videos were recorded between 09:00 and 16:00 at 23°C. *D. elegans* virgin females were used in all courtship assays in case any *D. elegans* female cues were necessary to elicit male wing display behavior. After each assay, both the male and female were aspirated back into an M food vial and left for up to 5 days, after which each male was frozen in individual 1.5 mL Eppendorf tubes for wing spot quantification (see Quantification of wing spots), genomic DNA (gDNA) extraction, and sequencing (see Library preparation and sequencing). All courtship videos (∼900 total) are available here: https://deepblue.lib.umich.edu/data/concern/data_sets/j098zb17n?locale=en.

### Quantification of wing display behavior

F_1_ hybrid and recombinant males from both backcross directions performed variable wing display behaviors during courtship as described previously (Yeh *et al.*, 2006; Yeh and True, 2014). To generate quantitative measurements of wing display variation between individuals, each courtship video was played using QuickTime (version 10.4) (Apple Inc., Cupertino, CA) software in a MacOS environment and digital screenshots were manually taken for each wing display bout, defined as a bilateral wing extension performed near the female (Supplementary Figure S1). Next, for each individual fly, wing display screenshots were compared to each other to identify the maximum wing display bout per fly, defined by comparing the distance between the tips of each wing relative to the center of the fly. These maximum wing display screenshots were then imported into ImageJ software (version 1.50i) (Wayne Rasband, National Institutes of Health, USA; http://rsbweb.nih.gov/ij/) to manually measure the “Maximum wing display angle” for F_1_ hybrid and recombinant males. In ImageJ, each screenshot image was inverted using the “Find Edges” function to enhance the contrast between the arena background and the edges of the fly wings (Supplementary Figure S1). Next, the “Polygon Selections” tool was used to fit an ellipse around the fly body using the “Fit Ellipse” function (Supplementary Figure S1). A Macros function (Supplementary File S1) was then used to generate major and minor axes inside the ellipse to identify the center of the fly body (Supplementary Figure S1). Finally, the “Angle Tool” was used to measure the “Maximum wing display angle” centering the vertex at the intersection of the major and minor axes and extended from wing tip to wing tip (Supplementary Figure S1). “Maximum wing display angle” varied between ∼50° and ∼220° between backcross recombinant individuals.

### Quantification of wing spots

Since wing spots fully form ∼24 h after eclosion in *D. elegans*, all parental male *D. elegans*, *D. gunungcola*, F_1_ hybrids, and backcross recombinants were aged at least 7 days before being frozen at −20C in 1.5 mL Eppendorf tubes. Next, using a 20 Gauge stainless steel syringe tip (Techcon) (cat# TE720100PK) the right wing of each fly was cut away from the thorax and placed on a glass microscope slide (Fisherbrand) (cat# 12-550-15) to image using either a Leica MZFLIII stereoscope equipped with a Leica DC480 microscope camera or a Canon EOS Rebel T6 camera equipped with a Canon MP-E 65 mm macro lens. Each camera was calibrated using an OMAX 0.1 mm slide micrometer to define pixel density in ImageJ software. JPEG images of wings were imported into ImageJ to measure wing spot size relative to total wing area (wing spot size / total wing area). Total wing area (wing length x wing width) was approximated using length and width proxies following methods described in Yeh and True (2014). Using the “Polygon Selections” tool, the margins of black pigmentation defining each “Wing spot size” was traced and the polygon area quantified in mm^2^ using the “Measure” function. “Wing spot size” varied between 0 mm^2^ (spotless) and 0.15 mm^2^ between recombinant individuals.

### Library preparation and sequencing

We estimated chromosome ancestry “genotypes” for 724 *D. elegans* backcross progeny and 241 *D. gunungcola* backcross progeny with a single Multiplexed Shotgun Genotyping (MSG) (Andolfatto *et al*., 2011) library using 965 barcoded adaptors following methods described in Cande *et al*., (2012). In brief, to extract gDNA from all male backcross individuals, single flies were placed into individual wells of 96-well (Corning, cat# 3879) plates containing a single steel grinding bead in each well (Qiagen, cat# 69989). Eleven plates in total were prepared for 965 individual gDNA extractions. gDNA was isolated and purified using the solid tissue extraction procedure from a Quick-DNA 96 Kit (Zymo, cat# D3012) and a paint shaker to homogenize tissue. gDNA was tagmented using a hyperactive version of Tn5 transposase charged with annealed adaptor oligos following the methods described in Picelli *et al*. (2014). Unique barcoded adaptor sequences were ligated to each sample of tagmented gDNA with 14 cycles of PCR using OneTaq 2x Master Mix (NEB, cat# M0482S), and all samples were pooled into a single multiplexed sequencing library. Agencourt AMPure XP beads (Beckman Coulter, cat# A63881) were used to size select ∼150-800 bp fragments and eluted in 35 uL of molecular grade water (Corning, cat# MT46000CI). The library was quantified by qPCR and sequenced in a single lane of Illumina HiSeq by the Janelia Quantitative Genomics Team.

In addition to generating the backcross sequencing library, both *D. elegans HK* and *D. gunungcola SK* parental species were sequenced at 20x coverage using an Illumina MiSeq Reagent Kit (v.3, 600 cycle PE) to facilitate genome assembly. In brief, gDNA was extracted using a Quick-DNA Microprep Kit (Zymo, cat# D4074) from 10 pooled females for each species and quantified on a Qubit 2.0 (Invitrogen). These samples were sent to the University of Michigan DNA Sequencing Core to prepare 300 bp paired-end libraries, which were quantified by qPCR and sequenced in a single lane of Illumina MiSeq.

### Genome assembly

In brief, Illumina reads from all 965 backcross recombinants were used to perform MSG on the Baylor College of Medicine *D. elegans* genome assembly (accession number: GCA_000224195.2). Using custom scripts in R and Python (https://github.com/masseyj/elegans), the recombination fraction between the Baylor and MSG contigs was calculated and plotted to manually tabulate joins and splits between newly assembled contigs. These new contigs were then used to assemble approximately chromosome length scaffolds in *D. elegans* (accession number: PRJNA590036) and partially assembled chromosomes in *D. gunungcola* (accession number: PRJNA590037).

### Marker generation with Multiplexed Shotgun Genotyping

Following methods described previously (Andolfatto *et al*., 2011; Cande *et al*., 2012), we used the MSG software pipeline (https://github.com/JaneliaSciComp/msg/tree/master/instructions) to perform data parsing and chromosome ancestry estimation to generate markers for quantitative trait locus (QTL) analysis. In brief, using data from the Illumina backcross sequencing library (see Supplementary File S2 for the number of reads per individual), we mapped reads to the assembled *D. elegans* and *D. gunungcola* parental genomes to estimate chromosome ancestry for each backcross individual. We generated 3,425 and 3,121 markers for the *D. elegans* and *D. gunungcola* backcrosses, respectively (Supplementary Files S3, S4), for QTL analysis. PDFs of chromosomal breakpoints for each recombinant are available here: https://deepblue.lib.umich.edu/data/concern/data_sets/j098zb17n?locale=en.

### QTL analysis

QTL analysis was performed using R/qtl (Broman *et al*., 2003) in R for Mac version 3.3.3 (R Core Team 2018) in a MacOS environment. Ancestry data for both backcross directions were imported into R/qtl using a custom script (https://github.com/dstern/read_cross_msg), which directly imports the conditional probability estimates produced by the Hidden Markov Model (HMM) of MSG (Andolfatto *et al*., 2011). We performed genome scans with a single QTL model using the “scanone” function of R/qtl and Haley-Knott regression (Haley and Knott, 1992) for “Wing spot size” and “Maximum wing display angle”. Note, for “Wing spot size”, 68 and 42 recombinants from the *D. elegans* and *D. gunungcola* backcross populations, respectively, were excluded from the QTL mapping because their wings were too damaged to quantify spot variation. Similarly, for “Maximum wing display angle”, 314 and 94 recombinants were excluded from the QTL mapping because these males did not perform any courtship behavior during the assay. Significance of QTL peaks at α = 0.01 was determined by performing 1000 permutations of the data. Effect sizes for each QTL peak were individually estimated by comparing the mean “Wing spot size” or “Maximum wing display angle” between individuals that inherited either *D. elegans* or *D. gunungcola* alleles at each QTL peak position.

Since we detected multiple QTL peaks on separate chromosomes for “Maximum wing display angle”, we tested for the presence of epistatic interactions using two methods: First, we performed two- and three-way ANOVAs comparing the effect of each QTL peak in multiple QTL peak genetic backgrounds and found no evidence of an interaction. For two-way ANOVAs, we tested for any statistically significant interactions for max wing display angles between two different QTL peaks in the *D. elegans* backcross. For three-way ANOVAs, we tested for any statistically significant interactions for max wing display angles between three different QTL peaks in the *D. gunungcola* backcross. Second, we performed genome-wide pairwise tests using the “scantwo” function of R/qtl and Haley-Knott regression to test for non-additive interactions across all markers; LOD significance thresholds at α = 0.05, 0.01, and 0.001 were determined by performing 1000 permutations of the data for each model (Supplementary Figure S2, Supplementary Tables S1,S2).

### Annotating the wing spot QTL interval

To annotate genes within the ∼440 Kbp fine-mapped wing spot locus, we performed nucleotide BLAST (BLASTn) (Johnson *et al*., 2008) searches against the *D. melanogaster* genome (taxid: 7227) using ∼10 Kbp windows of assembled *D. elegans* chromosomal regions spanning the wing spot QTL interval. Using the “GBrowse” tool on Flybase (Thurmond *et al*., 2018), we mapped regions of microsynteny to identify the orientation of each gene and exported the respective *D. melanogaster* coding region (CDS) FASTA sequences to align with the *D. elegans* X chromosome.

### In situ hybridization

Fly genomic DNA (gDNA) was extracted from ten homogenized *D. elegans* and *D. gunungcola* females using a Quick-DNA Microprep Kit (Zymo, cat# D3021). The following forward and reverse primers were designed and synthesized by Integrated DNA Technologies (IDT) to PCR amplify 321 bp DNA templates targeting exon 5 of the *omb* locus in *D. elegans*: 5’-GCTGAGGATCCATTCGCTAGATTTG-3’ and 5’-GTTGTTGGAACTAGAGTTGTTGGTG-3’, and *D. gunungcola*: 5’-GCTGAGGATCCATTCGCTAGATTTG-3’ and 5’-GTTGTTGGAACTGGAGTTGTTGGTG-3’. Reverse primers were designed beginning with a T7 RNA polymerase binding sequence (TAATACGACTCACTATAG) to facilitate *in vitro* transcription. Raw PCR products were then used to generate digoxigenin-labeled RNA probes using a T7 RNA *in vitro* transcription kit (Promega / Life Technologies). RNA was ethanol precipitated and resuspended in water to analyze on a Nanodrop. Each probe was stored at −20°C in 50% formamide before *in situ* hybridization.

All tissues underwent primary dissection in PBS, fixed for 30 mins in 4% PFA, washed 3X in PBT and underwent secondary dissection in PBT, were then washed 2X in MeOH, and 2X in EtOH before being stored at −20C. Male *D. elegans* and *D. gunungcola* L3 wing discs were dissected first to validate that our *omb* probes detected an mRNA expression pattern similar to *D. melanogaster* (Grimm and Pflugfelder, 1996; Supplementary Figure S3). Next, pupal wings were dissected at 30 and 48 h after pupal formation (APF) to probe for *omb* mRNA. To prepare pupal wings, appropriately staged pupae underwent a primary dissection: were cut in half along the anterior-posterior axis using Astra Platinum Double Edge Razor Blades, and fat body was washed out of the pupal casing using a pipette and PBS prior to fixation. After fixation, pupal wings underwent a secondary dissection to pull off the cuticle surrounding each wing and then washed using the procedure described above. Finally, *in situ* hybridization was carried out as previously described (Vincent *et al*., 2019). Briefly, we used an InsituPro VSi robot to rehydrate in PBT, fix in PBT with 4% PFA, and prehybridize in hybridization buffer for 1 hr at 65°C. Samples were then incubated with probe for 16 h at 65°C before washing with hybridization buffer and PBT. Samples were blocked in PBT with 1% bovine serum albumin (PBT+BSA) for 2 hours. Samples were then incubated with anti-digoxigenin Fab fragments conjugated to alkaline phosphatase (Roche) diluted 1:6000 in PBT+BSA. After additional washes, color reactions were performed by incubating samples with NBT and BCIP (Promega) until purple stain could be detected under a dissecting microscope. Samples were mounted in glycerol on microscope slides coated with poly-L-lysine and imaged at 10X magnification on a Leica DFC450C camera.

### Generating advanced recombinant introgressions on the X chromosome

To try to isolate the QTL effects for “Wing spot size” and “Maximum wing display angle” localized to the X chromosome according to the *D. elegans* backcross experiment, F_1_ hybrid females were generated using the procedures described above. F_1_ hybrid females were then backcrossed to *D. elegans* males, and backcross males lacking wing spots were isolated to measure “Maximum wing display angles” during courtship as described above. This procedure was repeated for seven generations to generate BC3-BC9 backcross individuals: backcross females were backcrossed en masse to *D. elegans* males, and BC3 backcross males lacking wing spots were isolated to measure “Maximum wing display angles” during courtship with *D. elegans* virgins (and so on to BC9). At each generation, an attempt was made to create stable introgression lines of advanced recombinant males lacking wing spots, but all failed to produce offspring, suggesting that *D. gunungcola* X-linked loci might also contain hybrid sterility factors. After seven generations of backcrossing, gDNA from all backcross males lacking wing spots was extracted and sequenced for MSG as described above. Backcross males lacking wing spots from BC4-BC9 were homozygous for *D. elegans* genomic regions across all autosomes but varied for the amount of *D. gunungcola* genome regions on the X chromosome.

### Introgression of black body color alleles from *D. gunungcola* into D. elegans

In the *D. gunungcola* backcross, QTL mapping for wing spot size revealed QTL peaks linked to Muller Element C and E when spotless recombinants were excluded from the analysis (Supplementary Figure S4; Supplementary Table S3). The Muller Element E QTL peak is located near the *ebony* gene, which appears to contribute to variation in body color between *D. elegans* and *D. gunungcola* (unpublished data). We therefore reasoned that introgressing dark body color from *D. gunungcola* into *D. elegans* would introgress the Muller Element E QTL peak underlying wing spot size differences. After six generations of backcrossing dark brown female recombinants with *D. elegans* males, we crossed dark brown male and female recombinants together to create black offspring homozygous for the introgressed region. We then performed MSG on a single, dark black introgression line and found that it was homozygous for ∼1.5 Mb of *D. gunungcola* alleles linked near the Muller Element E QTL peak (Supplementary Figure S4A,B).

### Observing and collecting wild *D. gunungcola* in Indonesia

Throughout early July 2018, *D. elegans* and *D. gunungcola* were recorded performing courtship in East Java, Indonesia on *Brugmansia sp.* flowers using Canon VIXIA HF R500 camcorders mounted to Manfrotto (MKCOMPACTACN-BK) aluminum tripods. Both species were observed in sympatry on flowers near Coban Rondo Waterfall in Batu, Batu City, East Java, Indonesia (−7.884985, 112.477311). After observing courtship, males and females were captured using a mouth pipette and gently aspirated into glass vials containing standard fly media (glucose, corn meal, yeast extract, and agar). Isofemale lines of *D. gunungcola* from Bumiaji District (Batu City, East Java Province, Indonesia) were established in the laboratory on standard fly media at 24°C temperature.

### Statistics

Statistical tests were performed in R for Mac version 3.3.3 (R Core Team 2018) using Student’s t-test (two-tailed) to test for statistically significant effects of pairwise comparisons of continuous data with normally distributed error terms. For tests comparing more than two groups, ANOVAs were performed with *post hoc* Tukey HSD for pairwise comparisons adjusted for multiple comparisons. See “QTL analysis” methods for statistical tests used during QTL mapping.

## Results and Discussion

### X-linked sequence divergence contributed to wing spot and wing display divergence

*D. elegans* males perform elaborate wing display dances (Video 1) in front of females during courtship, displaying the presence of darkly pigmented wing spots (Figure 1B), whereas its sibling species, *D. gunungcola*, lacks wing spots (Yeh *et al.*, 2006; Prud’homme *et al*., 2006) and wing displays (Figure 1B; Video 2). Despite these differences in sexual traits, *D. elegans* and *D. gunungcola* can mate and form viable F_1_ hybrids in the lab (Yeh *et al*., 2006; Yeh and True, 2014). Sequence divergence on the X chromosome has previously been implicated in the divergence of wing spots and wing display behavior (Yeh *et al*., 2006; Yeh and True, 2014). To confirm this effect of the X-chromosome, we quantified variation in wing spot size and wing display behavior in F_1_ hybrid males from reciprocal crosses between *D. elegans* and *D. gungungcola*. These F_1_ hybrids inherited their X chromosome from either *D. elegans* or *D. gunungcola* (whichever species was their mother) and autosomes from both species. Consistent with prior work, F_1_ hybrid males inheriting the X chromosome from *D. elegans* mothers (F_1_E) possessed wing spots, whereas F_1_ hybrid males inheriting the X chromosome from *D. gunungcola* mothers (F_1_G) did not (Figure 1C,D). These wing spots of F_1_E males were smaller, however, than the wing spots seen in *D. elegans* (Figure 1D, test, P = 0.02). Differences in wing display behavior were also apparent between F_1_E (Video 3) and F_1_G hybrids (Video 4), which is also consistent with prior work (Yeh *et al*., 2006; Yeh and True, 2014). More specifically, we found that although both F_1_ hybrids performed wing displays during courtship, F_1_E hybrids tended to open their wings more widely than F_1_G hybrids during display performance (Figure 1C). We quantified variation in this wing display trait between F_1_ hybrids by measuring the maximum bilateral wing display angles (Figure 1C) during courtship (see Methods). We found that F_1_E hybrids performed wing displays comparable to *D. elegans* males (Figure 1E, post-hoc Tukey HSD, P = 0.6), whereas F_1_G males showed, on average, lower display angles (Figure 1E, post-hoc Tukey HSD, P = 7.1 x 10^-5^). Together these data confirm that divergence of one or more loci on the X chromosome contribute to divergence in wing spot size and wing display behavior between *D. elegans* and *D. gunungcola*.

### Evolution of at least three loci contribute to wing spot divergence

To identify the location of X-linked (as well as autosomal) loci contributing to divergence in wing spot size, we quantified wing spot size variation in 656 recombinant males produced by backcrossing F_1_ hybrid females to *D. elegans* males and 199 recombinant males produced by backcrossing F_1_ hybrid females to *D. gunungcola* males. These backcross males showed a range of wing spot sizes (Figure 2A). Using Multiplexed Shotgun Genotyping (MSG) (Andolfatto *et al*., 2011), we inferred the allele most likely inherited from the F_1_ mother (*D. elegans* or *D. gunungcola*) for each genomic position in each recombinant. We then performed quantitative trait locus (QTL) mapping for wing spot size and identified a single, highly significant QTL peak on the X chromosome (Figure 2B and Table 1). In both backcross directions, variation linked to this wing spot QTL peak explained almost all of the difference in wing spot size between *D. elegans* and *D. gunungcola* (Figure 2C). Repeating the QTL mapping after excluding recombinant individuals lacking wing spots, however, allowed us to identify additional QTLs of smaller effect on Muller Elements C (chromosome 2R in *D. melanogaster*) and E (chromosome 3R in *D. melanogaster*) in the *D. gunungcola* (but not *D. elegans*) backcross population (Supplementary Figure S4A; Supplementary Table S3). Observing these QTL only in the *D. gunungcola* backcross populations suggests that they are caused by recessive *D. gunungcola* alleles, which are never homozygous in the *D. elegans* backcross population. Introgressing the QTL region on Muller Element E from *D. gunungcola* into *D. elegans* through 5 generations of backcrossing (Supplementary Figure S4C) reduced the size of wing spots (Supplementary Figure S4D, E). This region includes the *ebony* gene, which has previously been shown to be able to inhibit the development of dark pigments in *D. melanogaster* (Wittkopp *et al.*, 2002b). Crossing this introgression line to *D. elegans* masked most of the reduction in spot size (Supplementary Figure 4D, E), consistent with the *D. gunungcola* QTL allele being recessive to the *D. elegans* allele. Taken together, these data indicate that the majority of wing spot divergence between *D. elegans* and *D. gunungcola* maps to a single, large-effect QTL on the X chromosome, but that wing spot size is also influenced by loci on Muller Elements C and E.

**Figure 2.**
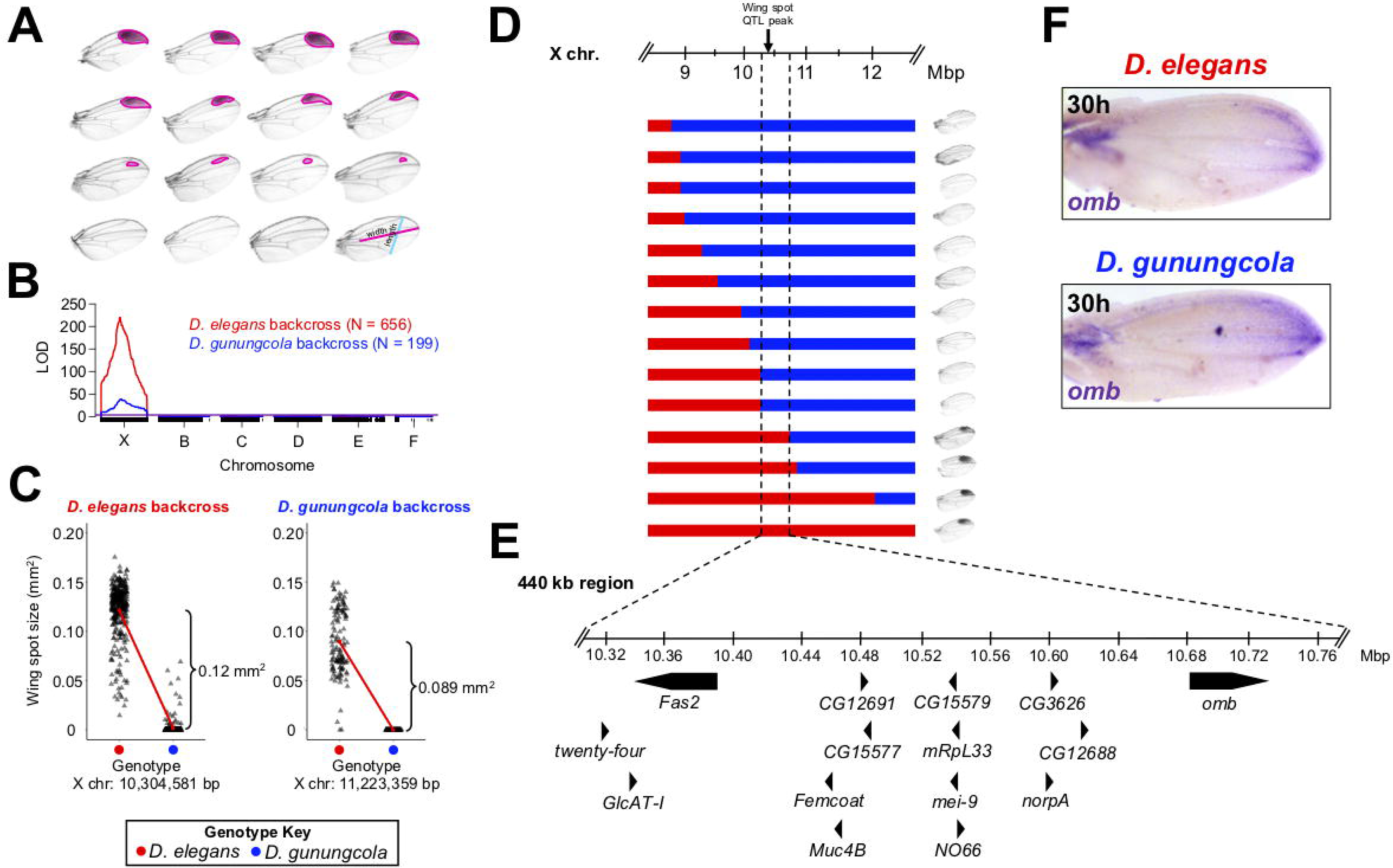
QTL analysis, effect plots, and *in situ* hybridization for wing pigmentation divergence. (A) Wing spots vary in size and shape in *D. elegans* and *D. gunungcola* backcross recombinants. Wing spots were traced (pink) and quantified relative to proxies for total wing area (length x width) using ImageJ software (see Methods). (B) Wing spot QTL map for the *D. elegans* (red) and *D. gunungcola* (blue) backcross. LOD (logarithm of the odds) is indicated on the y-axis. The x-axis represents the physical map of Muller Elements X, B, C, D, E, and F based on the *D. elegans* assembled genome (see Methods). While *D. elegans* and *D. gunungcola* have six separate chromosomes (Yeh *et al*., 2006; Yeh and True, 2014), they are each syntenic with the *D. melanogaster* genome accordingly: X = X, B = 2L, C = 2R, D = 3L, E = 3R, F = 4. Individual SNP markers are indicated with black tick marks along the x-axis. Horizontal red and blue lines mark p = 0.01 for the *D. elegans* and *D. gunungcola* backcross, respectively. (C) Effect plots for the X chromosome QTL peak from the *D. elegans* backcross (left) and *D. gunungcola* backcross (right). (D) *D. elegans* and *D. gunungcola* backcross recombinants containing X chromosome breakpoints immediately flanking the wing spot QTL peak were aligned to compare the effects of each on wing pigmentation. Regions in red represent *D. elegans* linked loci, and regions in blue represent *D. gunungcola* linked loci. Recombinants possessing *D. elegans* loci to the left of ∼10.32 Mbp are spotless, while recombinants possessing *D. elegans* loci to the right of ∼10.74 Mbp possess dark wing spots. (E) Two recombinants define the wing spot locus to a ∼440 Kbp region containing 15 candidate genes. *omb* is the strongest wing pigmentation candidate gene given evidence from prior work (see Results and Discussion). (F) *In situ* hybridization of *D. elegans* and *D. gunungcola* pupal wings probed for *omb* mRNA (purple) at 30 h after pupal formation (APF) (see Supplementary Figure S8 for additional replicates). Gray triangles represent individual replicates.

**Table 1.**
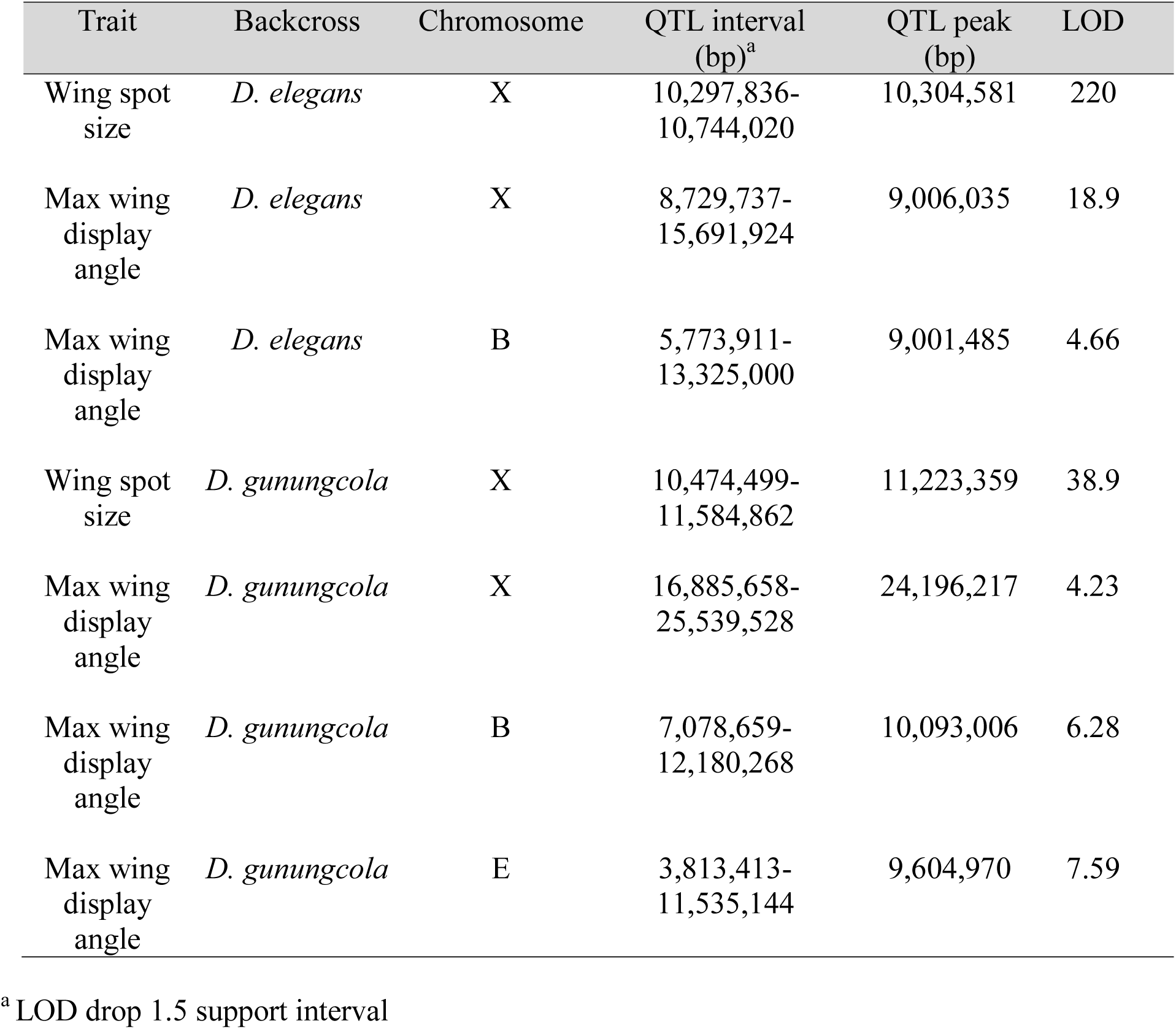
QTLs detected for wing spot size and maximum wing display angle divergence.

### A 440 kb locus behaves like a genetic switch for wing spots

To further refine the X-linked QTL, we more closely examined the genotypes and phenotypes of recombinants with inferred crossover positions immediately flanking the wing spot QTL peak (Figure 2D, Supplementary Figure S5). Doing so allowed us to identify a ∼440 kb region containing a QTL that acts like a genetic switch controlling the presence or absence of the wing spot (Figure 2D, Supplementary Figure S5). This region includes 15 genes (Figure 2E) and notably excludes the X-linked pigmentation gene, *yellow*, which has previously been suggested to contribute to wing spot development and evolution (Wittkopp *et al*., 2002a; Gompel *et al*., 2005; Prud’homme *et al*., 2006; Yeh *et al*., 2006; Arnoult *et al*., 2013; Yeh and True, 2014; Supplementary Figure 6). One of these 15 genes is *optomotor-blind* (*omb*) (Figure 2E), which encodes a T-box-containing transcription factor (Pflugfelder *et al*., 1992a; Pflugfelder *et al*., 1992b) that has previously been implicated in pigmentation patterning (Thompson, 1959; Kopp and Duncan, 1997), pigmentation evolution (Brisson *et al*., 2004), and distal wing patterning (Grim and Pflugfelder, 1996). In *D. melanogaster*, gain- and loss-of-function *omb* alleles cause expansion and contraction of abdominal pigmentation bands, respectively (Kopp and Duncan, 1997), and variation in abdominal pigmentation in *D. polymorpha* is strongly associated with polymorphisms at the *omb* locus (Brisson *et al*., 2004).

Although we identified two nonsynonymous protein coding changes between *D. elegans* and *D. gunungcola* (Supplementary File S5), *omb* is required for the development of many structures throughout the body (Pflugfelder, 2009); we, therefore, reasoned that genetic divergence in *omb* would be more likely to affect its expression than its protein function (Stern and Orgogozo, 2008). To look for differences in *omb* expression between *D. elegans* and *D. gunungcola* that might affect wing spot development, we used *in situ* hybridization to detect *omb* mRNA in the developing wing of both species (Figure 2F). In *D. melanogaster*, *omb* is expressed in a broad stripe that overlaps the wing pouch region in larval L3 wing discs (Grimm and Pflugfelder, 1996). *omb* expression in the wing pouch is required for distal wing development, as demonstrated by *D. melanogaster omb* hypomorphs that show disrupted distal wing tip development in adults (Grimm and Pflugfelder, 1996). We hypothesized, therefore, that differences in *D. elegans* and *D. gunungcola omb* expression patterning during pupal wing development might prefigure changes in wing spot pigmentation observed in adult males, similar to the changes in *wingless* expression shown to prefigure wing spots in *D. guttifera* (Werner *et al.*, 2010). Consistent with the expression of *omb-lacZ* in pupal wings of *D. melanogaster* (Álamo Rodríguez et al., 2004), we detected *omb* mRNA in the wing hinge and distal wing tip 30 h after puparium formation (APF) in *D. elegans* and *D. gunungcola* (Figure 2F). We were unable to identify any consistent differences in the *omb* expression patterns between *D. elegans* and *D. gunungcola* although it is possible that w e may not have detected subtle differences in expression patterns. In addition, it is possible that the changes in *omb* protein sequence contribute to differences in wing spot patterning, or that other genes in the minimal mapped interval are the true cause of the difference in wing spot patterning.

### Evolution at multiple loci contributed to wing display divergence

To identify loci contributing to divergence in wing display behavior, we quantified variation in maximum wing display angles (see Methods) in 410 *D. elegans* and 147 *D. gunungcola* backcross recombinant males, again observing a range of phenotypes (Figure 3A). We identified multiple significant QTL contributing to variation in wing display (Figure 3B; Table 1). In the *D. elegans* backcross, we mapped a QTL on the X chromosome that co-localized with the wing spot QTL (Figure 3B,E; Table 1). We also mapped a QTL on Muller Element B (chromosome 2L in *D. melanogaster*) (Figure 3B; Table 1). In the *D. gunungcola* backcross, we mapped QTLs on the X chromosome as well as Muller Elements B and E (Figure 2E; Table 1). These differences in QTL peaks mapped using backcrosses to *D. elegans* and *D. gunungcola* suggest that *D. elegans* and *D. gunungcola* alleles affecting wing display behavior are recessive and/or interact epistatically with divergent sites elsewhere in the genome.

**Figure 3.**
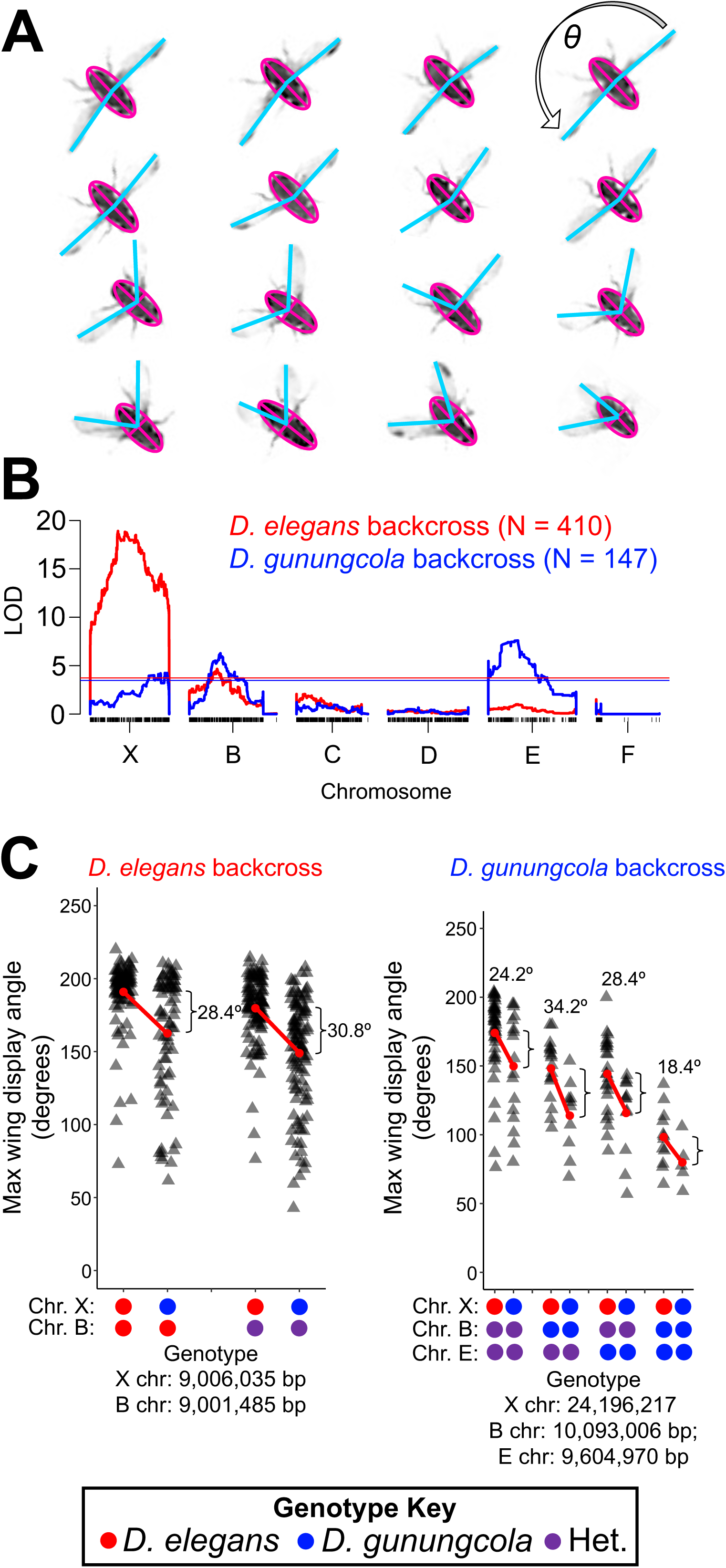
QTL analysis and effect plots for wing display divergence. (A) Maximum wing display angles varied in *D. elegans* and *D. gunungcola* backcross recombinants. Maximum wing display angles were quantified by measuring the angle between each wing tip using ImageJ software (see Methods). (B) Maximum wing display QTL map for the *D. elegans* (red) and *D. gunungcola* (blue) backcross. LOD is indicated on the y-axis. Individual SNP markers are indicated with black tick marks along the x-axis. Horizontal red and blue lines mark P = 0.01 for the *D. elegans* and *D. gunungcola* backcross, respectively. (C) Effect plots for the X chromosome and Muller Element B QTL peaks from the *D. elegans* backcross (left) and for the X, Muller Element B, and E QTL peaks from the *D. gunungcola* backcross (right). No epistatic interactions were detected between QTLs (see Methods) (Two-way ANOVA: F_1,402_ = 0.146; P = 0.70 for the *D. elegans* backcross; Three-way ANOVA: F_1,137_ = 0.050 (X:B), 0.034 (X:E), 1.75 (B:E), 0.799 (X:B:E); P = 0.82 (X:B), 0.86 (X:E), 0.19 (B:E), 0.37 (X:B:E) for the *D. gunungcola* backcross). Gray triangles represent individual replicates.

To test for epistatic interactions contributing to wing display divergence, we performed a two-dimensional genome scan to search for non-additive interactions across all markers in both backcross directions and found no significant interactions (Supplementary Figure S2; Supplementary Tables S1,S2). We also tested for evidence of non-additive interactions among the wing display QTL peaks themselves by performing two- and three-way ANOVAs in the *D. elegans* and *D. gunungcola* backcrosses, respectively, and found no evidence of significant interactions between loci (Figure 3C). Instead, each wing display QTL peak appears to behave approximately additively, with *D. gunungcola* alleles contributing to lower maximum wing display angles (Figure 3C). Surprisingly, the effect of the X-linked QTL on wing display angle in the *D. gunungcola* backcross in multiple genetic backgrounds was similar to the estimated effect size of the X-linked QTL in the *D. elegans* backcross (compare panels in Figure 3C) despite the much lower LOD score of the X-linked QTL in the *D. gunungcola* backcross population (Figure 3B; Table 1). We suggest that while the detected QTL in the *D. gunungcola* backcross appear to interact additively with each other, undetected QTL elsewhere in the genome are likely masking the X-effect in the *D. gunungcola* backcross map. While the purpose of the two-dimensional genome scan (Supplementary Figure S2; Supplementary Tables S1,S2) was to detect these effects, our sample size is likely too small to identify small-effect epistatic interactions.

### Males lacking wing spots perform normal wing displays

While it remains unclear which gene evolved to cause the majority of wing spot divergence, fine-mapping the locus controlling the presence or absence of the wing spot allowed us to test whether the locus that turns off wing spots in *D. gunungcola* also affects wing display behavior. To perform this test, we introgressed *D. gunungcola* alleles causing a loss of the wing spot into *D. elegans* by repeated backcrossing (see Methods). We recovered three introgression lines lacking wing spots and found that all three lines had inherited the ∼440 kb region observed in mapping experiments to act like a genetic switch controlling wing spot development (Figure 4A,B), independently confirming the causal role of the switch region in wing spot divergence. We noticed, however, that several advanced recombinants developed a wing spot “shadow” (Figure 4B), possibly due to the effects of other *D. elegans* alleles affecting wing spot development. We next asked whether the spotless advanced recombinants performed wing displays with lower wing display angles than *D. elegans* males. Surprisingly, we found that all advanced recombinants inheriting the *D. gunungcola* allele eliminating the wing spot performed wing displays indistinguishable from *D. elegans* males during courtship (Figure 4B,C; Videos 5-7). Thus, the loci controlling the wing spot and courtship behavior are genetically separable.

**Figure 4.**
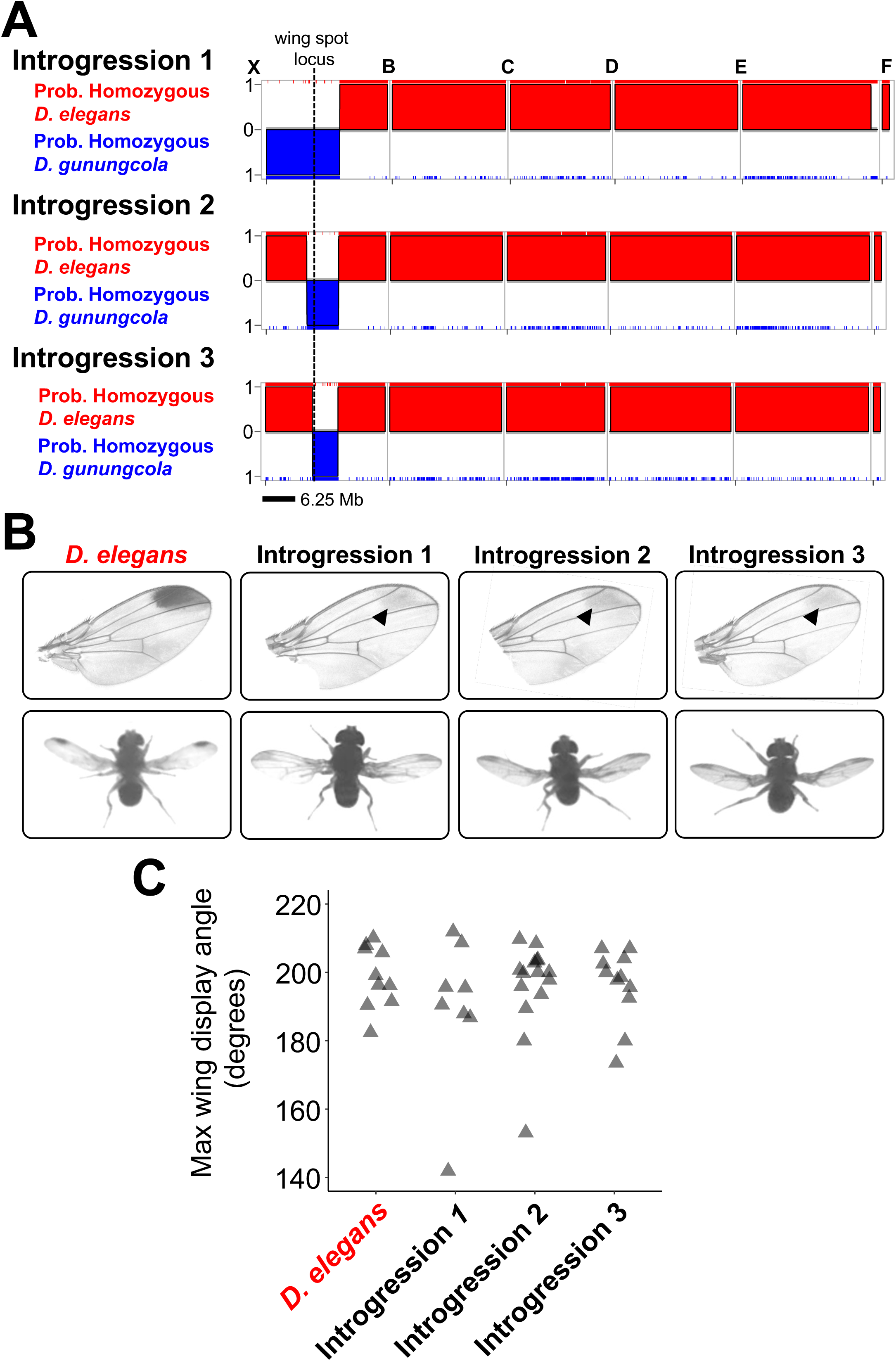
*D. elegans* males possessing the *D. gunungcola* wing spot locus perform normal wing displays. (A) Multiplexed Shotgun Genotyping (MSG) (Andolfatto *et al*., 2011) was used to estimate genome-wide ancestry assignments for three introgression lines generated by repeatedly backcrossing the *D. gunungcola* wing spot QTL region into a *D. elegans* genetic background (see Methods). The posterior probability that a region is homozygous for *D. elegans* (red) or *D. gunungcola* (blue) ancestry is plotted along the y-axis. The dotted line marks the location of the fine-mapped wing spot region (Figure 2D,E; Table 1). (B). None of the introgressions possessed dark wing spots (although a light wing spot “shadow” is visible). (B,C) Every introgression performed max wing display angles indistinguishable from *D. elegans* males (One-way ANOVA: F_3,42_ = 0.449; P = 0.72). Gray triangles represent individual replicates.

The repeated co-evolution of male-specific wing spots and wing display behavior in multiple species (Kopp and True, 2002) combined with the presence of overlapping QTL for these traits on the X chromosome (Yeh *et al*., 2006; Yeh and True, 2014; and this study) suggested that a single pleiotropic gene might be contributing to the evolution of both traits. The finding that *D. elegans* introgression lines lacking a wing spot performed a normal wing display argues against this hypothesis and indicates instead that these two traits arose independently between this species pair. To further investigate how these divergent traits might have evolved, we observed courtship behavior in a wild population of *D. gunungcola* in Indonesia; to the best of our knowledge, all prior studies of *D. gunungcola* pigmentation and courtship used the one previously available lab strain (Sultana *et al*., 1999). Surprisingly, we found that all *D. gunungcola* males observed in the wild population sampled lacked wing spots (Supplementary Figure 7) but performed wing displays (Videos 8,9), confirming that these are genetically distinct traits. The wing displays performed by these flies appeared to show a lower maximum wing extension angle than *D. elegans* (Videos 1,10), similar to the wing display behavior seen in F1 hybrids between *D. elegans* and *D. gunungcola* with *D. gunungcola* mothers (Video 4). Analysis of new lab strains founded by flies captured from this *D. gunungcola* population showed similar male courtship behavior in the lab as observed on flowers (Video 11). We therefore conclude that although the absence of wing spots appears fixed in *D. gunungcola*, the absence of wing display behavior does not. It remains to be seen whether the lack of wing display in the strain collected in 1999 resulted from polymorphic alleles segregating within *D. gunungcola* or a change that occurred since this strain was brought into the laboratory. Assuming that the loss of the wing spot and wing display behavior are derived traits in *D. gunungcola* (Prud’homme *et al*., 2006), these observations suggest that the loss of male-specific wing spots predates the loss of male wing display behavior in this species.

## Conclusions

Male-specific wing spots and wing display behavior have co-evolved in *Drosophila* multiple times (Kopp and True, 2002). By studying the genetic basis of these divergent traits between *D. elegans* and *D. gunungcola*, we showed that the changes in wing spot and wing display were not caused by changes in a single, pleiotropic gene despite overlapping QTL (Yeh *et al*., 2006; Yeh and True, 2014). Rather, we found that distinct loci contribute to divergence in each of these traits, with the genetic architecture of divergent wing behavior being more complex than that of the divergent wing spot pigmentation. Both traits were affected by divergent gene(s) located on the X chromosome that are in physical linkage, however, causing alleles of these distinct loci to be co-inherited. This linkage might have facilitated the coordinated evolution of these traits.

The specific genes contributing to divergence in wing spot and wing display remain unknown, but *optomotor-blind* is a strong candidate for the X-linked gene contributing to the loss of the wing spot. Introgression lines and additional sampling of *D. gunungcola* from a wild population also showed that the loss of wing spots and wing display are not inexorably linked: in both cases, males lacking wing spots still performed a wing display behavior. Coordinated evolution of morphological and behavioral traits such as these is often observed in animal species, but it is often unclear which change evolved first. In this case at least, it seems that the divergence of morphology preceded the divergence of behavior.

## Acknowledgements

We thank members of the Wittkopp, Stern, and Rebeiz labs for helpful discussions. For fly strains, we thank John True (Stony Brook University). For guidance throughout the *in situ* hybridization work, we thank Mark Rebeiz (University of Pittsburgh). For arranging the Material Transfer Agreement for *D. gunungcola* and *D. elegans*, we thank Nia Kurniawan (University of Brawijaya, Indonesia); for hosting us in Indonesia, we thank Karuniawan Wicaksono (University of Brawijaya, Indonesia); for assistance with field collections, we thank Hagus Tarno (University of Brawijaya, Indonesia). Funding: University of Michigan, Department of Ecology and Evolutionary Biology, Peter Olaus Okkelberg Research Award, National Institutes of Health (NIH) training grant T32GM007544, and Howard Hughes Medical Institute Janelia Graduate Research Fellowship to J.H.M.; NIH R01 GM089736 and 1R35GM118073 to PJW.

## Author Contributions

Jonathan H. Massey, Conceptualization, Data curation, Formal analysis, Funding acquisition, Validation, Investigation, Visualization, Methodology, Writing—original draft, Writing—review and editing; Gavin R. Rice, Formal analysis, Validation, Investigation, Methodology, Writing— review and editing; Anggun Firdaus, Investigation, Methodology, Writing—review and editing; Chi-Yang Chen, Investigation, Methodology; Shu-Dan Yeh, Funding acquisition, Investigation, Methodology; David L. Stern, Supervision, Funding acquisition, Conceptualization, Data curation, Formal analysis, Investigation, Visualization, Writing—original draft, Project administration, Writing—review and editing; Patricia J. Wittkopp, Supervision, Funding acquisition, Conceptualization, Data curation, Formal analysis, Investigation, Visualization, Writing—original draft, Project administration, Writing—review and editing.

---

**Supplementary Figure S1 ImageJ procedure for measuring maximum wing display angles**

Screenshots of each wing display were captured for every recombinant courtship video. The maximum wing display bout was identified for each fly by quickly comparing screenshots that varied in wing display angles (from wing tip to wing tip) and picking by eye the display with the largest angle. Next, for each fly, the maximum wing display angle was quantified in ImageJ by using 1) Find Edges function, 2) polygon tool to Fit Ellipse around the fly body, 3) Ellipse Macros (Supplementary File S1) to fit the major and minor axes of the ellipse, and 4) draw Angle tool, fitting the angle vertex at the major and minor axes intersection to calculate the wing display angle from wing tip to wing tip.

**Supplementary Figure S2 LOD scores estimated from a two-dimensional, two QTL scan of maximum wing display angles**

(A) For the *D. elegans* backcross, the Interaction LODi, which estimates the likelihood that the effect of genotypes at one marker depend on genotypes at another, is displayed in the upper left triangle; the Full LODf, which estimates the effect of both additive and non-additive interactions between genotypes (see Supplementary Table S2 for LOD thresholds), is displayed in the lower right triangle (Broman *et al*., 2003). The color scale on the right indicates LOD values for LODi (left) and LODf (right). (B) For the *D. gunungcola* backcross, the Interaction LODi is displayed in the upper left triangle; the Full LODf (see Supplementary Table S3 for LOD thresholds) is displayed in the lower right triangle. The color scale on the right indicates LOD values for LODi (left) and LODf (right).

**Supplementary Figure S3 *In situ* hybridization of *D. elegans* and *D. gunungcola* L3 wing discs**

Male *D. elegans* (left) and *D. gunungcola* (right) L3 wing discs were dissected and stained with probes targeting *omb* mRNA.

**Supplementary Figure S4 Effects of Muller Element E on wing spot divergence**

(A) Wing spot QTL map for *D. elegans* (red) and *D. gunungcola* (blue) backcross recombinants. Note, all recombinant individuals that lacked wing spots were removed from this QTL analysis to identify loci contributing to wing spot size variation independent of wing spot presence or absence. LOD (logarithm of the odds) is indicated on the y-axis. The x-axis represents the physical map of Muller Elements X, B, C, D, E, and F based on the *D. elegans* assembled genome (see Methods). Individual SNP markers are indicated with black tick marks along the x-axis. Horizontal red and blue lines mark P = 0.01 for the *D. elegans* and *D. gunungcola* backcross, respectively. (B) Images illustrating *D. elegans* and *D. gunungcola* body color differences. (C) Multiplexed Shotgun Genotyping (MSG) (Andolfatto *et al*., 2011) was used to estimate genome-wide ancestry assignments for a single introgression line generated by repeatedly backcrossing *D. gunungcola* into a *D. elegans* genetic background (see Methods). The posterior probability that a region is homozygous for *D. elegans* (red) or *D. gunungcola* (blue) ancestry is plotted along the y-axis. (D) Representative wing spot images of *D. elegans* and *D. gunungcola* species parents, the introgression line genotyped in (B), and an F_1_ heterozygote generated by crossing *D. elegans* females to introgression males. (E) Quantification of wing spot size differences between each genotype. Results of Tukey HSD *post hoc* tests following one-way ANOVA are shown (One-way ANOVA F_2,88_ = 78.6; P < 2.0 x 10^-16^; post-hoc Tukey HSD was significant between *D. elegans* and Introgression: P < 1.0 x 10^-7^, *D. elegans* and F1 Inrogression/*D. elegans* heterozygote: P = 0.02, and Introgression and Inrogression/*D. elegans* heterozygote: P < 1.0 x 10^-7^. Gray triangles represent individual replicates.

**Supplementary Figure S5 Fine-mapping the wing spot locus**

*D. elegans* and *D. gunungcola* backcross recombinants containing X chromosome breakpoints immediately flanking the wing spot QTL peak were aligned to compare the effects of each on wing pigmentation. Regions in red represent *D. elegans* linked loci, and regions in blue represent *D. gunungcola* linked loci. All recombinants possessing *D. gunungcola* loci to the right of ∼10.95 Mbp are spotless.

**Supplementary Figure S6 Effects of the *yellow* gene on wing spot size and wing display behavior in *D. elegans***

(A) Loss-of-function *D. elegans HK yellow* mutants develop smaller wing spots than *D. elegans HK* wild-type males (Student’s t-test; t = 4.7759; df = 15.28; P = 0.0002; two-tailed) and (B) show lower maximum wing display angles (Student’s t-test; t = 3.0294; df = 50.82; P = 0.004; two-tailed).

**Supplementary Figure S7 Male wings from new *D. gunungcola* isolates**

Male *D. gunungcola* from five newly collected isofemale lines in Indonesia do not develop wing spots.

**Supplementary Figure S8 *In situ* hybridization of *D. elegans* and *D. gunungcola* pupal wings probed for *omb* mRNA**

*omb* mRNA (purple) was probed at 30 and 48 h after pupal formation (APF) in both males and females.

## Videos

**Video 1 *D. elegans HK* wing display behavior**

**Video 2 *D. gunungcola SK* courtship and copulation**

**Video 3 F_1_E wing display behavior**

**Video 4 F_1_G wing display behavior**

**Video 5 Introgression 1 wing display behavior**

**Video 6 Introgression 2 wing display behavior**

**Video 7 Introgression 3 wing display behavior**

**Video 8 *D. gunungcola* wing display behavior at Coban Rondo Waterfall in East Java, Indonesia (Version 1)**

**Video 9 *D. gunungcola* wing display behavior at Coban Rondo Waterfall in East Java, Indonesia (Version 2)**

**Video 10 *D. elegans* wing display behavior in Tumpang, Indonesia**

**Video 11 *D. gunungcola* (Batu City, Indonesia) wing display behavior in the laboratory**

