## Supplementary figures and images for "Co-evolving wing spots and mating displays are genetically separable traits in *Drosophila*"

### Supplementary Figure S1

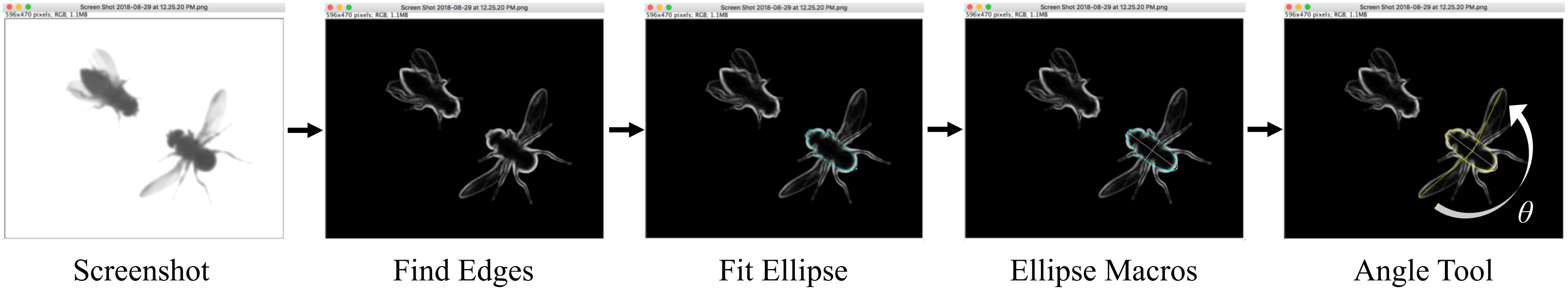

### Supplementary Figure S2

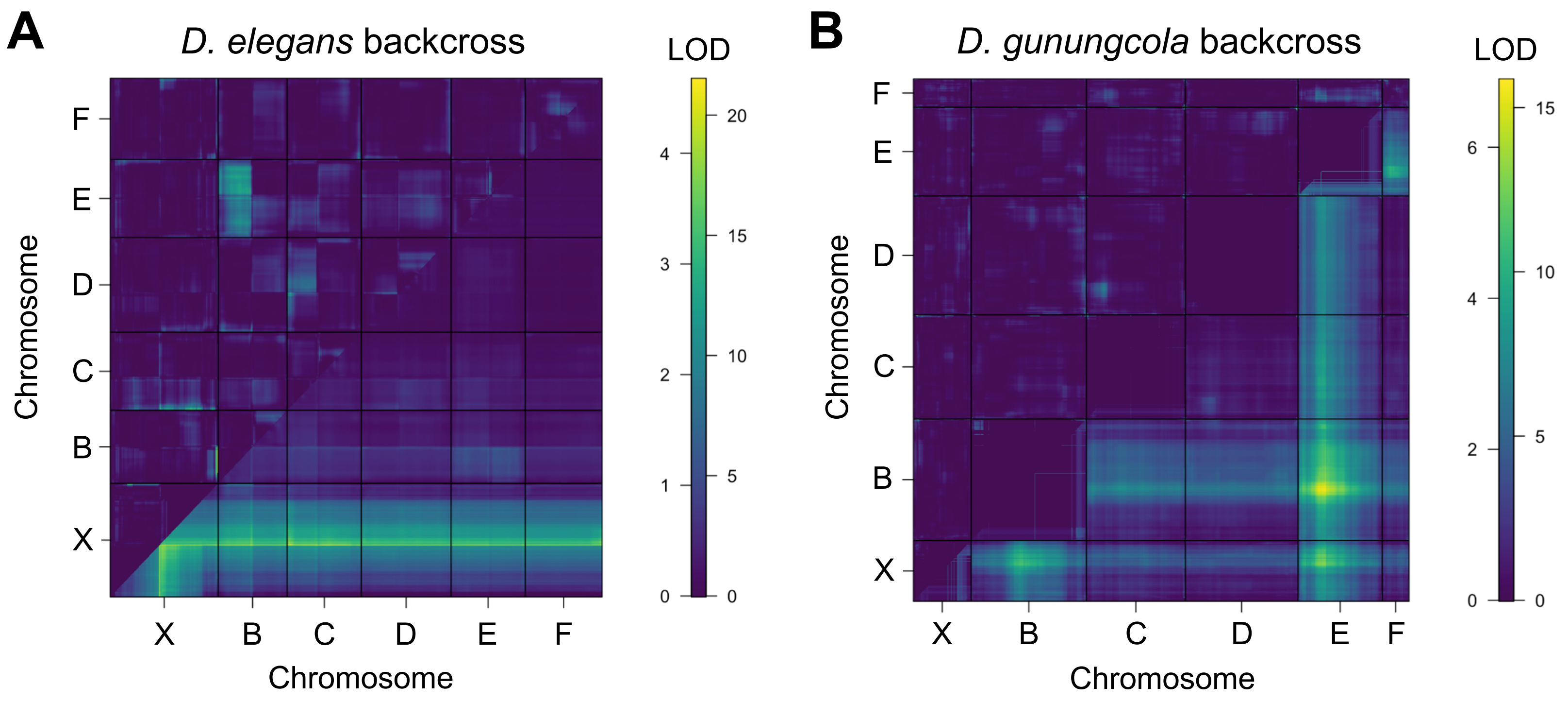

### Supplementary Figure S3

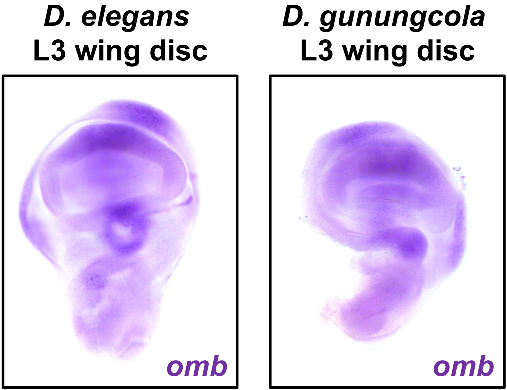

### Supplementary Figure S4

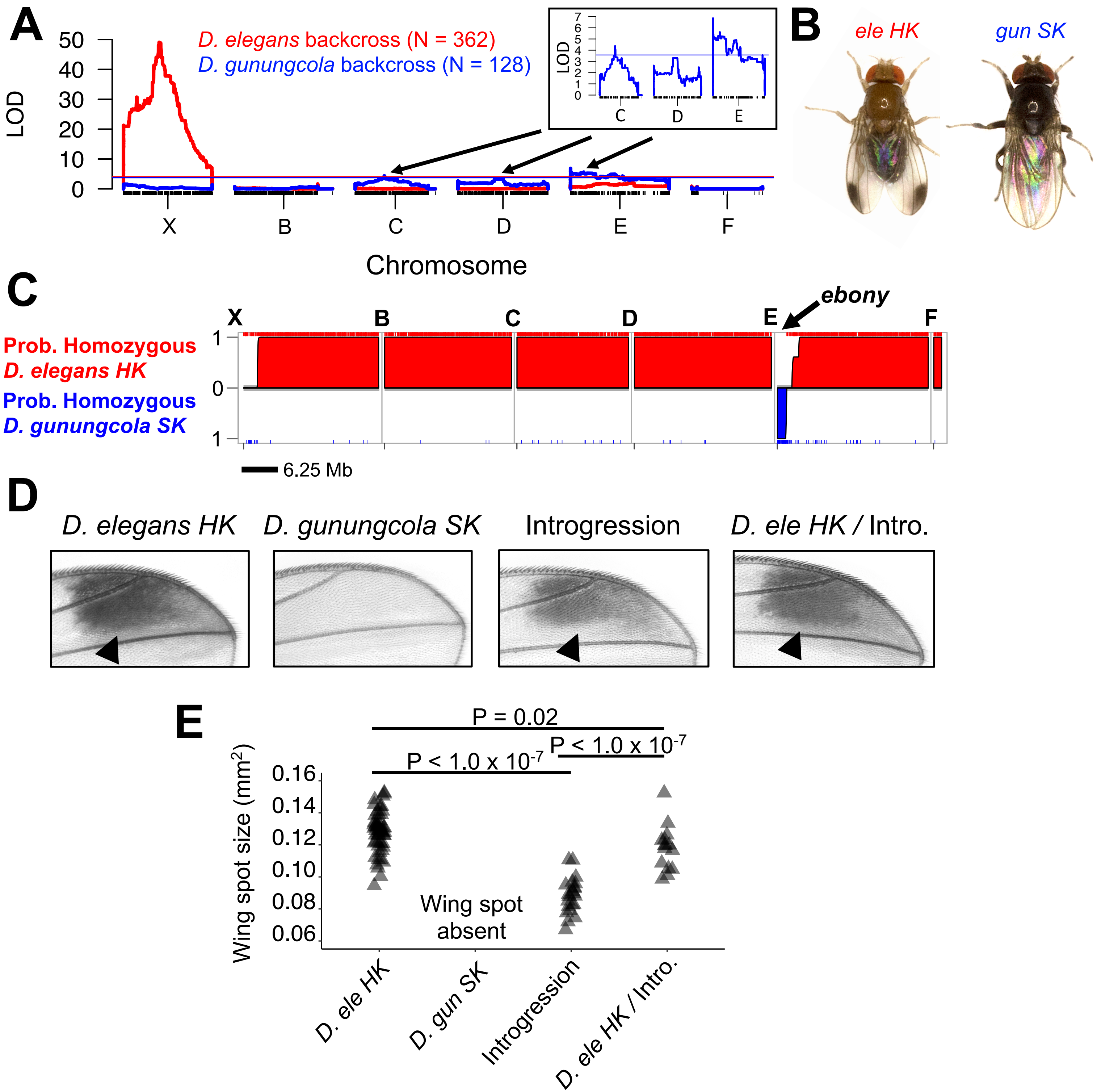

### Supplementary Figure S5

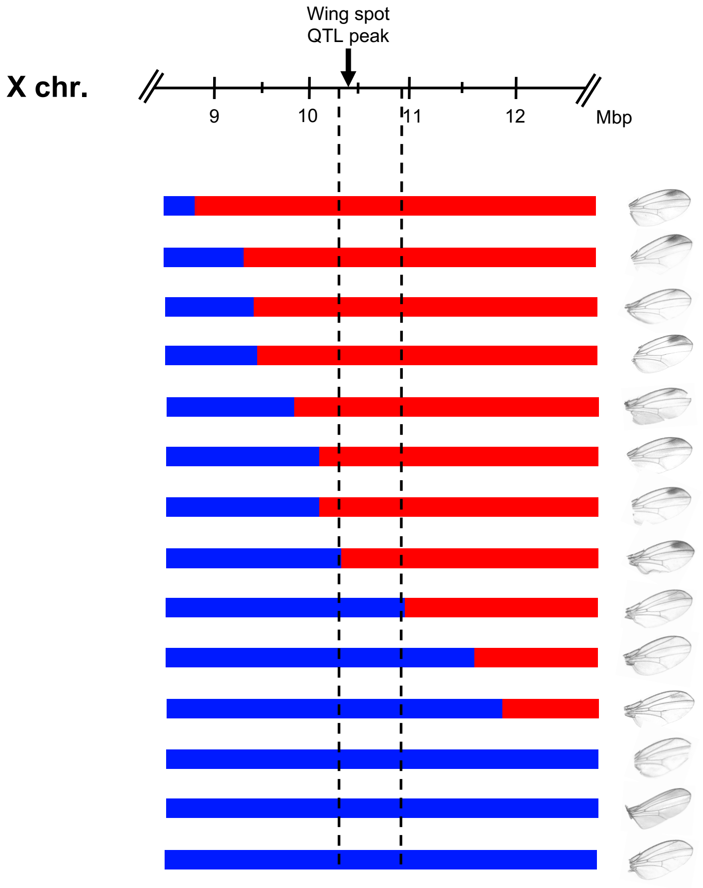

### Supplementary Figure S6

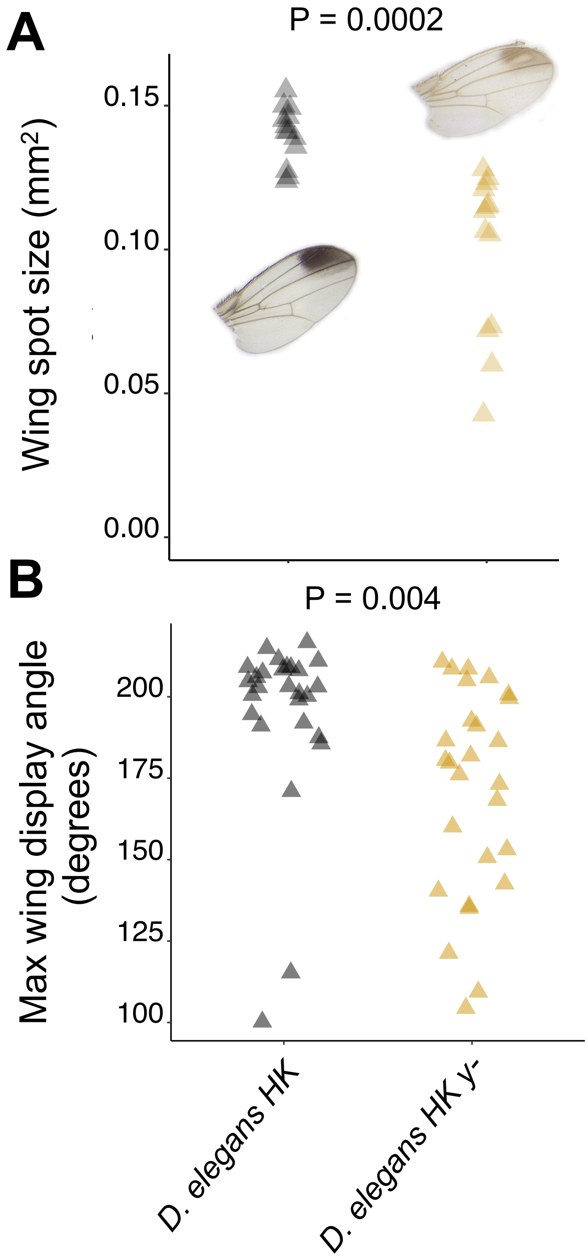

### Supplementary Figure S7

♂ *D. gunungcola* C62

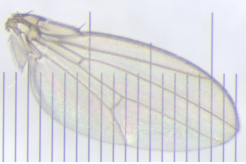

♂ *D. gunungcola* C89

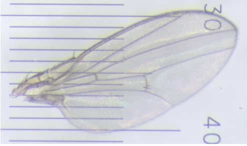


*D. gunungcola* C91

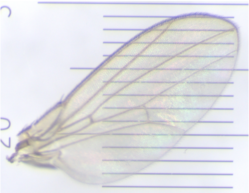

*D. gunungcola* C95

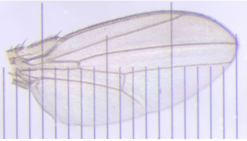

*D. gunungcola* C99

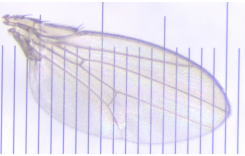

### Supplementary Figure S8

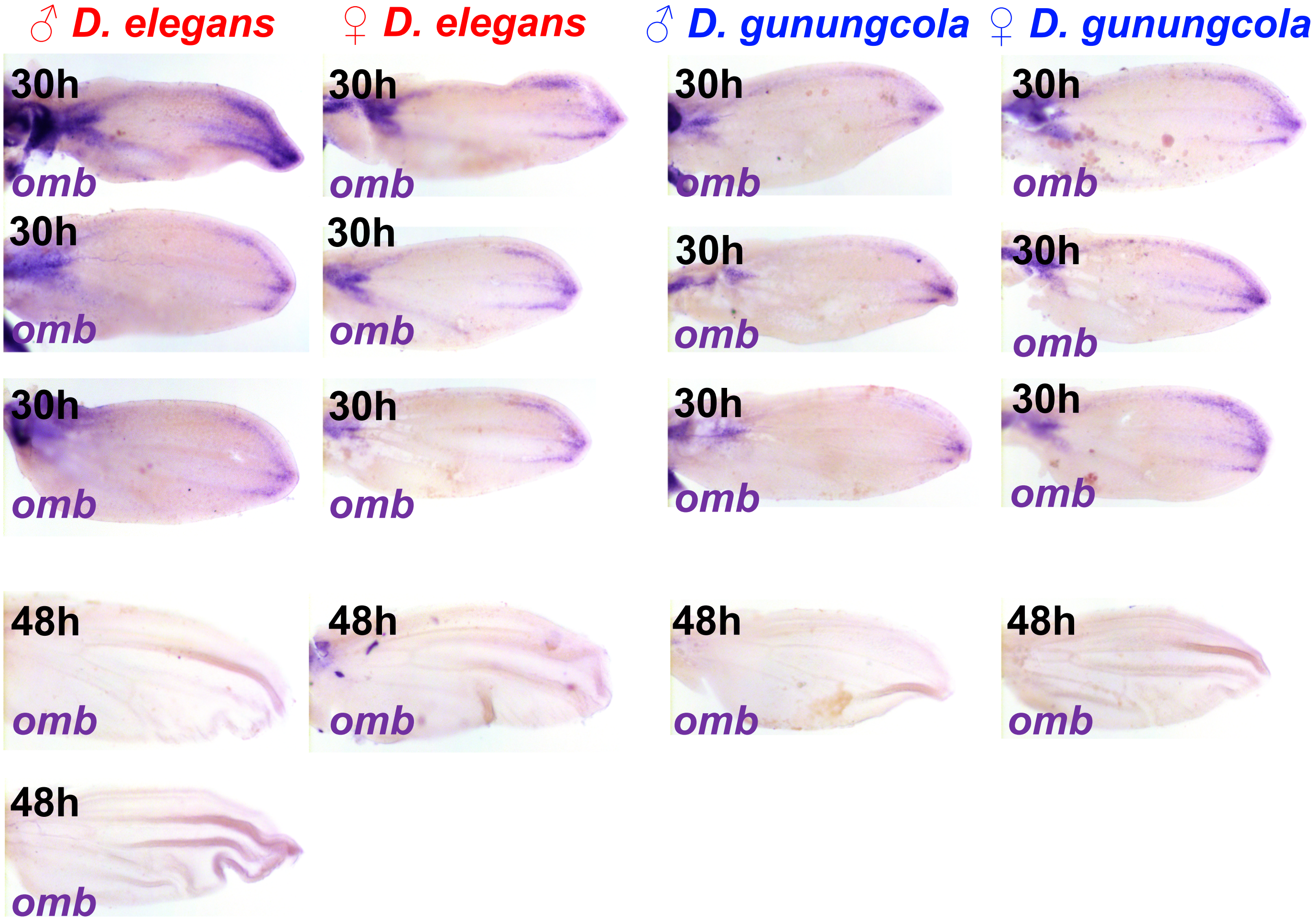
