## Supplementary File S5 for "Co-evolving wing spots and mating displays are genetically separable traits in *Drosophila*"

!Title Dgun_Dele_omb_CDS;protein

!Format DataType=nucleotide;protein

#Dgun_omb_CDS

ATGAGATACGACGTCCAGGAGCTGCTACTTCATCAGTCTGCTGAGGATCCATTCGCTAGATTTGCCAATGGGATGGCATATCATCCATTTCTGCAGCTAACGCAACGACCCACTGACTTCAGCGTATCCTCGCTGTTGACGGCGGGTAGTAATACTAATAATAGTAATAATAGTAATAACAACAGCAGTGGAAACACCAACTCCAACAAYAATTCCAACAACACCAATAATAACACCAACAATCTTGTGGCCGTTTCACCAACGGGTGGTGGTGCGCAGTTATCGCCGCAGAGCAACCACAGCAGCAGCAACACCACCACCACCAGCAACACCAACAACTCCAGTTCCAACAACAACAATAATAATAATAAYAACAACAATAATAATAATAATAAYAACAACCATACAAATAATAATAATAAYACGAATCAAAAACAAGGACACCACTTGAGCACCACCGAAGAAACAACATCGCCAGCTGGGACACCACCACCACCGCCCACAATTGTTGGTGTACCGTCAATACCACCGCCCAACAACAGCAGCAGCAACAACAACACAGCTGCCCATCCGTCGCACCACCCAACAACAACAGCTGCAGCCAGGGAAGCTCATCACTCACCAAATACGGGTGCCGCTGCACCACCCACTGGCGTAACGGGTCTACCGCCACCCACACCGCCGCACCACCTCCAGCAACAACAGCAGCACCAGCAGCAGGCACCACCACCACCACCGCCCTACTTTCCCGCTGCGGCACTGGCCGCTCTAGCTGGCAGTCCGGCCGGACCGCATCCGGGCCTCTATCCCGGCGGTGGCCTACGCTTTCCGCCCCACCACCCCGGTGCCCACCCACACGCCCACCACCTGGGCAGCGCCTACACCACCGCCGAGGACGTCGTCCTTGCCTCGGCCGTAGCCCACCAGCTGCATCCGGCGATGCGACCGCTGCGGGCCCTGCAGCCCGAGGACGACGGGGTCGTCGACGATCCCAAGGTCACGCTGGAGGGCAAGGACCTGTGGGAGAAGTTCCACAAGCTGGGCACCGAAATGGTCATCACCAAGAGCGGCAGACAAATGTTTCCGCAAATGAAATTTCGTGTTTCGGGACTGGATGCCAAGGCTAAATACATCTTGCTACTGGACATCGTGGCGGCGGACGATTATCGTTATAAATTTCATAATAGTCGCTGGATGGTGGCTGGGAAAGCGGACCCCGAGATGCCAAAACGCATGTATATCCATCCAGATTCGCCCACAACGGGTGAGCAATGGATGCAGAAAGTTGTTTCATTTCACAAATTAAAATTGACCAACAATATTAGTGATAAACATGGATTTACGATCCTGAACTCGATGCACAAGTACCAGCCGCGTTTCCACCTGGTGCGAGCCAATGACATCCTGAAGCTGCCGTACTCCACGTTTCGCACGTACGTCTTCAAGGAGACCGAGTTCATCGCCGTCACTGCATATCAAAATGAGAAGATAACTCAATTGAAAATCGATAACAATCCCTTCGCGAAGGGCTTTCGTGATACTGGTGCTGGCAAGCGGGAAAAGAATTGTTACAGGCAGGCGTTGATGTCGAACCGAGGGTCCGATTCGGACAAGCTAAACCCCACGCATGTTAGCAGCTCGCGGGCACCGCTCCACCTGGGCCACGCCGGTCGTCCGCCCCACCTGCACCCCCATGCCGGATTGCTGGACAACCAGCAGGACGACGACGACAAGCTCCTGGACGTTGTGGGTCCGCCGCAGAGTCCGCTCCTGCCGCTCAGCCACTCGCTGCAGCAGATGCACGCACACCAGCACTCCGCCTTGGCCGCCTGGTTCAATCACCTGGCCGGAGCGGGAGCCGGCGCCTCGGAGCACGCCGCAGCGGCGGCCGCCAATGCCAGTGCGGAGGATGCACTGCGTCGCCGCCTGCAGGCGGATGCGGACGCGGAGCGCGATGGCAGCGACTCGAGCTGCTCGGAGAGCGTCGGCGGCAGCACCGGCGGTGCCTTTAGGCCCACCTCGACGGGCAGTCCCAAGGAGGCGGTGGTGGGCGCGGCTGCCGCAGCGGCTGCCCTCAATGCCGGCGGCGGTGGCGGCGGCAGCTACCCATCGCCGAATATATCGGTGGGTCCGCCGATCCACCCGTCGCCGCACCTGTTGCCTTACCTCTATCCCCACGGCCTCTATCCGCCGCCGCACCTGGGCCTGCTCCACAATCCCGCAGCGGCAGCGGCCATGAGTCCGGCCGGCCTCAATCCCGGCCTGCTCTTCAATGCCCAGCTGGCGCTGGCCGCCCAGCATCCGGCCCTGTTTGGCCACGCCTACGCGGCGGCGGGACACACGCCGGTCTCGCCRCTGCAGGGCTTGAAGAGCCACCGCTTCTCGCCGTACAGCTTGCCGGGCAGCCTGGGCTCCGCCTTTGATGCAGTCACRCCCGGCTCGAATGCCAATCGCTCGGGTGACCCGCCCGGAATTCCAGCCGTGGAGAATGGCCCCCGGAGCTTGAGCTCCAGTCCGCGACCTCGGCCCGCCTCCCACTCGCCGCCCACGCGGCCCATCTCCATGTCACCCACCACACCACCTTCGCTGATGAAGCGTGGCGCTGGCGGTGGCGGCGGCGGTGGCGGYGGCGGTGGCGGTGGGGTGTCCCAATCCCAGCACTCCCCTTCGGAACTCAAGAGCATGGAGAAGATGGTCAATGGGCTGGAGGTGCAGCACAATGGTAGTGCGGCGGCGGCGGCAGCAGCACTTCAACTGGCCGAGGAGGCTGCTCAGCACCACCATCACACCACCCAGGAGCACCACTCAGTGCATGCGCATTCGCATTCGCATTCGCACCATCAGCAGCAGCAGCAGCAGCAGTCGCACCACCAGCAGCAGCAGCAGCAGTCACACCACCACCACCAGACGCACCTCCATTCGCAATCGCAATCGCATCACGGAGCGAGTGCGGATCAGTGA

#Dele_omb_CDS

ATGAGATACGACGTCCAGGAGCTGCTACTTCATCAGTCTGCTGAGGATCCATTCGCTAGATTTGCCAATGGGATGGCATATCATCCATTTCTGCAGCTAACGCAACGACCCACTGACTTCAGCGTATCCTCGCTGTTGACGGCGGGTAGTAATACTAATAATAGTAATAATAGTAACAACAACAGCAGTGGAAACACCAACTCGAACAACAATTCCAACAACACCAATAGTAACACCAACAATCTTGTGGCCGTTTCACCAACGGGTGGTGGTGCGCAGTTATCGCCGCAGAGCAACCACAGCAGCAGCAACACCACCACCACCAGCAACACCAACAACTCTAGTTCCAACAACAACAATAATAATAATAACAACAACAATAATAATAATAATAACAACAACCATACAAATAATAATAATAACACGAATCAAAAACAAGGACACCACTTGAGCACCACCGAAGAAACAACATCGCCAGCTGGGACACCACCACCACCGCCCACAATTGTTGGTGTACCGTCAATACCACCGCCCAACAACAGCAGCAGCAACAACAACACAGCTGCCCATCCGTCGCACCACCCAACAACAACAGCTGCAGCCAGGGAGGCTCATCACTCACCAAATACGGGTGCCGCTGCACCACCCACTGGCGTAACGGGTCTACCGCCACCCACACCGCCGCACCACCTCCAGCAACAACAGCAGCACCAGCAGCAGGCACCACCACCACCACCGCCCTACTTTCCCGCTGCGGCACTGGCCGCTCTAGCTGGCAGTCCGGCCGGACCGCATCCGGGCCTCTATCCCGGCGGTGGCCTACGCTTTCCGCCCCACCACCCCGGTGCCCACCCACACGCCCACCATCTGGGCAGCGCCTACACCACCGCCGAGGACGTCGTCCTTGCCTCGGCCGTCGCCCACCAGCTGCATCCGGCGATGCGACCGCTGCGGGCCCTGCAGCCCGAGGACGACGGGGTCGTCGACGATCCCAAGGTCACGCTGGAGGGCAAGGACCTGTGGGAGAAGTTCCACAAGCTGGGCACCGAAATGGTCATCACCAAGAGCGGCAGACAAATGTTTCCGCAAATGAAATTTCGTGTTTCGGGACTGGATGCCAAGGCTAAATACATCTTGCTACTGGACATCGTGGCGGCGGACGATTATCGTTATAAATTTCATAATAGTCGCTGGATGGTGGCTGGGAAAGCGGACCCCGAGATGCCAAAACGCATGTATATCCATCCAGATTCGCCCACAACGGGTGAGCAATGGATGCAGAAAGTTGTTTCATTTCACAAATTAAAATTGACCAACAATATTAGTGATAAACATGGATTTACGATCCTGAACTCGATGCACAAGTACCAGCCGCGTTTCCACCTGGTGCGAGCCAATGACATCCTGAAGCTGCCGTACTCCACGTTTCGCACGTACGTCTTCAAGGAGACCGAGTTCATCGCCGTCACTGCATATCAAAATGAGAAGATAACTCAATTGAAAATCGATAACAATCCCTTCGCAAAGGGCTTTCGTGATACTGGTGCTGGAAAGCGGGAAAAGAATTGTTACAGGCAGGCGTTGATGTCGAACCGAGGGTCCGATTCGGACAAGCTAAACCCCACGCATGTGAGCAGCTCGCGGGCACCGCTCCACCTGGGCCACGCCGGTCGTCCGCCCCATCTGCATCCCCATGCCGGATTGCTGGACAACCAGCAGGACGACGACGACAAGCTCCTGGACGTCGTGGGTCCGCCGCAGAGTCCGCTCCTGCCGCTCAGCCACTCGCTGCAGCAGATGCACGCCCACCAGCACTCCGCCTTGGCCGCCTGGTTCAATCACCTGGCCGGAGCGGGAGCCGGCGCCTCGGAGCACGCCGCAGCGGCGGCCGCCAATGCCAGTGCGGAGGATGCACTGCGTCGCCGCCTGCAGGCGGATGCGGACGCGGAGCGCGATGGCAGCGACTCGAGCTGCTCGGAGAGCGTCGGCGGGAGCACCGGCGGTGCCTTTAGGCCCACCTCGACGGGCAGTCCCAAGGAGGCGGTGGTGGGCGCGGCTGCCGCAGCGGCTGCCCTCAATGCCGGCGGCGGTGGCGGCGGCAGCTACCCATCGCCGAATATATCGGTGGGTCCGCCGATCCACCCGTCGCCGCACCTGTTGCCTTACCTCTATCCCCACGGCCTCTATCCGCCGCCGCACCTGGGCCTGCTCCACAATCCCGCAGCGGCAGCGGCCATGAGTCCGGCCGGCCTCAATCCCGGCCTGCTCTTCAATGCCCAGCTGGCGCTGGCCGCCCAGCATCCGGCCCTGTTTGGCCACGCCTACGCGGCGGCGGGACACACGCCGGTCTCGCCGCTGCAGGGCTTGAAGAGCCACCGCTTCTCGCCGTACAGCTTGCCGGGCAGCCTGGGCTCCGCCTTTGATGCAGTCACGCCCGGCTCGAATGCCAATCGCTCGGGTGACCCGCCCGGAATGCCAGCCGTGGAGAATGGCCCCCGGAGCTTGAGCTCCAGTCCGCGACCTCGGCCCGCCTCCCACTCGCCGCCCACGCGACCCATCTCCATGTCACCCACCACACCACCTTCGCTGATGAAGCGTGGCGCTGGCGGTGGCGGCGGCGGTGGCGGTGGCGGTGGCGGTGGGGTGTCCCAATCCCAGCACTCCCCTTCGGAACTCAAGAGCATGGAGAAGATGGTCAATGGGCTGGAGGTGCAGCACAATGGTAGTGCGGCGGCGGCGGCAGCAGCACTTCAACTGGCCGAGGAGGCTGCTCAGCACCACCATCACACCACCCAGGAGCACCACTCAGTGCATGCGCATTCGCATTCGCACTCGCACCATCAGCAGCAGCAGCAGCAGCAGTCGCACCACCAGCAGCAGCAGCAGCAGTCACACCACCACCACCAGACGCACCTCCATTCGCAATCGCAATCGCATCATGGAGCGAGTGCGGATCAGTGA

#Dgun_omb_protein

MRYDVQELLLHQSAEDPFARFANGMAYHPFLQLTQRPTDFSVSSLLTAGSNTNNSNNSNNNSSGNTNSNXNSNNTNNNTNNLVAVSPTGGGAQLSPQSNHSSSNTTTTSNTNNSSSNNNNNNNXNNNNNNNXNNHTNNNNXTNQKQGHHLSTTEETTSPAGTPPPPPTIVGVPSIPPPNNSSSNNNTAAHPSHHPTTTAAAREAHHSPNTGAAAPPTGVTGLPPPTPPHHLQQQQQHQQQAPPPPPPYFPAAALAALAGSPAGPHPGLYPGGGLRFPPHHPGAHPHAHHLGSAYTTAEDVVLASAVAHQLHPAMRPLRALQPEDDGVVDDPKVTLEGKDLWEKFHKLGTEMVITKSGRQMFPQMKFRVSGLDAKAKYILLLDIVAADDYRYKFHNSRWMVAGKADPEMPKRMYIHPDSPTTGEQWMQKVVSFHKLKLTNNISDKHGFTILNSMHKYQPRFHLVRANDILKLPYSTFRTYVFKETEFIAVTAYQNEKITQLKIDNNPFAKGFRDTGAGKREKNCYRQALMSNRGSDSDKLNPTHVSSSRAPLHLGHAGRPPHLHPHAGLLDNQQDDDDKLLDVVGPPQSPLLPLSHSLQQMHAHQHSALAAWFNHLAGAGAGASEHAAAAAANASAEDALRRRLQADADAERDGSDSSCSESVGGSTGGAFRPTSTGSPKEAVVGAAAAAAALNAGGGGGGSYPSPNISVGPPIHPSPHLLPYLYPHGLYPPPHLGLLHNPAAAAAMSPAGLNPGLLFNAQLALAAQHPALFGHAYAAAGHTPVSXLQGLKSHRFSPYSLPGSLGSAFDAVXPGSNANRSGDPPGIPAVENGPRSLSSSPRPRPASHSPPTRPISMSPTTPPSLMKRGAGGGGGGGXGGGGGVSQSQHSPSELKSMEKMVNGLEVQHNGSAAAAAAALQLAEEAAQHHHHTTQEHHSVHAHSHSHSHHQQQQQQQSHHQQQQQQSHHHHQTHLHSQSQSHHGASADQ

#Dele_omb_protein

MRYDVQELLLHQSAEDPFARFANGMAYHPFLQLTQRPTDFSVSSLLTAGSNTNNSNNSNNNSSGNTNSNNNSNNTNSNTNNLVAVSPTGGGAQLSPQSNHSSSNTTTTSNTNNSSSNNNNNNNNNNNNNNNNNNHTNNNNNTNQKQGHHLSTTEETTSPAGTPPPPPTIVGVPSIPPPNNSSSNNNTAAHPSHHPTTTAAAREAHHSPNTGAAAPPTGVTGLPPPTPPHHLQQQQQHQQQAPPPPPPYFPAAALAALAGSPAGPHPGLYPGGGLRFPPHHPGAHPHAHHLGSAYTTAEDVVLASAVAHQLHPAMRPLRALQPEDDGVVDDPKVTLEGKDLWEKFHKLGTEMVITKSGRQMFPQMKFRVSGLDAKAKYILLLDIVAADDYRYKFHNSRWMVAGKADPEMPKRMYIHPDSPTTGEQWMQKVVSFHKLKLTNNISDKHGFTILNSMHKYQPRFHLVRANDILKLPYSTFRTYVFKETEFIAVTAYQNEKITQLKIDNNPFAKGFRDTGAGKREKNCYRQALMSNRGSDSDKLNPTHVSSSRAPLHLGHAGRPPHLHPHAGLLDNQQDDDDKLLDVVGPPQSPLLPLSHSLQQMHAHQHSALAAWFNHLAGAGAGASEHAAAAAANASAEDALRRRLQADADAERDGSDSSCSESVGGSTGGAFRPTSTGSPKEAVVGAAAAAAALNAGGGGGGSYPSPNISVGPPIHPSPHLLPYLYPHGLYPPPHLGLLHNPAAAAAMSPAGLNPGLLFNAQLALAAQHPALFGHAYAAAGHTPVSPLQGLKSHRFSPYSLPGSLGSAFDAVTPGSNANRSGDPPGMPAVENGPRSLSSSPRPRPASHSPPTRPISMSPTTPPSLMKRGAGGGGGGGGGGGGGVSQSQHSPSELKSMEKMVNGLEVQHNGSAAAAAAALQLAEEAAQHHHHTTQEHHSVHAHSHSHSHHQQQQQQQSHHQQQQQQSHHHHQTHLHSQSQSHHGASADQ
