## Supplementary Table S1 for "Co-evolving wing spots and mating displays are genetically separable traits in *Drosophila*"

Supplementary Table S2. Results of two-QTL scan for max wing display angle in *D. elegans* backcross

| **Chromosomes** | **Full^a^** | **Two QTL^b^** | **Interaction^c^** | **Full Additive^d^** | **Two Additive^e^** |
| --- | --- | --- | --- | --- | --- |
| X:X | 20.02*** | 1.947 | 0.0144 | 20.006*** | 1.933 |
| X:B | 21.33*** | 3.253 | 0.0161 | 21.311*** | 3.237* |
| X:C | 21.49*** | 3.412 | 1.6614 | 19.824*** | 1.750 |
| X:D | 18.84*** | 0.763 | 0.1868 | 18.649*** | 0.576 |
| X:E | 19.36*** | 1.289 | 0.6103 | 18.752*** | 0.678 |
| X:F | 19.09*** | 1.016 | 0.1577 | 18.931*** | 0.858 |
| B:B | 4.71* | 0.788 | 0.2569 | 4.453* | 0.531 |
| B:C | 5.57** | 1.644 | 0.0926 | 5.474** | 1.551 |
| B:D | 6.14** | 2.216 | 1.5895 | 4.549* | 0.627 |
| B:E | 6.77** | 2.845 | 1.7483 | 5.019** | 1.096 |
| B:F | 5.01** | 1.083 | 0.7518 | 4.254* | 0.331 |
| C:C | 3.20 | 1.116 | 0.1116 | 3.091 | 1.004 |
| C:D | 4.49* | 2.405 | 1.8687 | 2.623 | 0.536 |
| C:E | 3.27 | 1.187 | 0.5495 | 2.724 | 0.637 |
| C:F | 2.46 | 0.371 | 0.0888 | 2.369 | 0.282 |
| D:D | 2.31 | 1.938 | 0.0584 | 2.250 | 1.880 |
| D:E | 2.04 | 1.255 | 0.8588 | 1.176 | 0.396 |
| D:F | 1.46 | 1.090 | 0.7354 | 0.725 | 0.355 |
| E:E | 4.82* | 4.044* | 1.4233 | 3.401 | 2.620 |
| E:F | 1.71 | 0.929 | 0.5999 | 1.109 | 0.329 |
| F:F | 3.13 | 2.798 | 0.7486 | 2.381 | 2.049 |

*P < 0.05, ** P < 0.01, *** P < 0.001

LOD significance thresholds at α = 0.05, 0.01, and 0.001 were determined by performing 1000 permutations of the data for each model

^a^ Maximum LOD score for the full model with interactions allowed (LOD thresholds: 4.2, 5.0, 6.5 for α = 0.05, 0.01, and 0.001, respectively)

^b^ Difference between the Full LOD and the maximum single-QTL LOD for the chromosome pair (LOD thresholds: 4.0, 4.5, 5.5 for α = 0.05, 0.01, and 0.001, respectively)

^c^ Difference between the maximum Full and Full Additive LODs (LOD thresholds: 3.9, 4.0, 4.7 for α = 0.05, 0.01, and 0.001, respectively)

^d^ Maximum LOD score for two QTLs with only additive interactions allowed (LOD thresholds: 4.4, 4.9, 5.75 for α = 0.05, 0.01, and 0.001, respectively)

^e^ Difference in LODs between the Full Additive model and the maximum single QTL model for the chromosome pair (LOD thresholds: 3.0, 3.5, 3.9 for α = 0.05, 0.01, and 0.001, respectively)
