## Supplementary Table S2 for "Co-evolving wing spots and mating displays are genetically separable traits in *Drosophila*"

Supplementary Table S3. Results of two-QTL scan for max wing display angle in *D. gunungcola* backcross

| **Chromosomes** | **Full^a^** | **Two QTL^b^** | **Interaction^c^** | **Full Additive^d^** | **Two Additive^e^** |
| --- | --- | --- | --- | --- | --- |
| X:X | 5.44** | 1.212 | 0.56884 | 4.87* | 0.643 |
| X:B | 10.75*** | 5.288** | 0.00381 | 10.74*** | 5.285*** |
| X:C | 5.65** | 1.423 | 0.00193 | 5.65** | 1.421 |
| X:D | 5.29* | 1.059 | 0.53058 | 4.76* | 0.529 |
| X:E | 13.10*** | 5.490** | 0.26490 | 12.83*** | 5.225*** |
| X:F | 5.60** | 1.373 | 0.39196 | 5.21** | 0.981 |
| B:B | 7.61*** | 2.152 | 1.23537 | 6.38*** | 0.917 |
| B:C | 6.97*** | 1.512 | 0.40271 | 6.57*** | 1.109 |
| B:D | 6.42*** | 0.958 | 0.68002 | 5.74** | 0.278 |
| B:E | 14.70*** | 7.098*** | 0.15974 | 14.54*** | 6.938*** |
| B:F | 6.48*** | 1.024 | 0.03732 | 6.45*** | 0.987 |
| C:C | 3.11 | 1.977 | 0.18173 | 2.93 | 1.795 |
| C:D | 2.82 | 1.694 | 1.37202 | 1.45 | 0.322 |
| C:E | 9.20** | 1.596 | 0.40867 | 8.79*** | 1.187 |
| C:F | 3.03 | 1.899 | 1.01448 | 2.01 | 0.884 |
| D:D | 2.25 | 1.907 | 0.61844 | 1.63 | 1.288 |
| D:E | 8.24*** | 0.630 | 0.02843 | 8.21*** | 0.601 |
| D:F | 2.29 | 1.386 | 0.98636 | 1.30 | 0.400 |
| E:E | 8.53*** | 0.922 | 0.50165 | 8.03*** | 0.421 |
| E:F | 10.84*** | 3.237 | 2.08462 | 8.76*** | 1.152 |
| F:F | 3.55 | 2.652 | 1.05307 | 2.50 | 1.599 |

*P < 0.05, ** P < 0.01, *** P < 0.001

LOD significance thresholds at α = 0.05, 0.01, and 0.001 were determined by performing 1000 permutations of the data for each model

^a^ Maximum LOD score for the full model with interactions allowed (LOD thresholds: 4.4, 5.3, 6.0 for α = 0.05, 0.01, and 0.001, respectively)

^b^ Difference between the Full LOD and the maximum single-QTL LOD for the chromosome pair

(LOD thresholds: 4.2, 4.8, 5.5 for α = 0.05, 0.01, and 0.001, respectively)

^c^ Difference between the maximum Full and Full Additive LODs (LOD thresholds: 4.0, 4.4, 4.9 for α = 0.05, 0.01, and 0.001, respectively)

^d^ Maximum LOD score for two QTLs with only additive interactions allowed LOD thresholds: 4.6, 5.1, 5.75 for α = 0.05, 0.01, and 0.001, respectively)

^e^ Difference in LODs between the Full Additive model and the maximum single QTL model for the chromosome pair (LOD thresholds: 3.2, 3.4, 4.0 for α = 0.05, 0.01, and 0.001, respectively)
