## Supplementary Table S3 for "Co-evolving wing spots and mating displays are genetically separable traits in *Drosophila*"

Supplementary Table S1 QTLs detected for wing spot size, excluding spotless individuals

| Trait | Backcross | Chromosome | QTL interval (bp)^a^ | QTL peak (bp) | LOD |
| --- | --- | --- | --- | --- | --- |
| Wing spot size | *D. elegans* | X | 10,117,675-10,748,234 | 10,303,766 | 49.1 |
| Wing spot  size | *D. gunungcola* | C | 6,655,757-12,279,025 | 8,420,192 | 4.37 |
| Wing spot  size | *D. gunungcola* | E | 10,907-4,009,870 | 12,292 | 6.85 |

^a^ LOD drop 1.5 support interval
